# Three times NO: no relationship between frontal alpha asymmetry and depressive disorders in a multiverse analysis of three studies

**DOI:** 10.1101/2020.06.30.180760

**Authors:** Aleksandra Kołodziej, Mikołaj Magnuski, Anastasia Ruban, Aneta Brzezicka

## Abstract

For decades, the frontal alpha asymmetry (FAA) - a disproportion in EEG alpha oscillations power between right and left frontal channels - has been one of the most popular measures of depressive disorders (DD) in electrophysiology studies. Patients with DD often manifest a left-sided FAA: relatively higher alpha power in the left versus right frontal lobe. Recently, however, multiple studies failed to confirm this effect, questioning its reproducibility. Our purpose is to thoroughly test the validity of FAA in depression by conducting a multiverse analysis - running many related analyses and testing the sensitivity of the effect to changes in the analytical approach - on data from three independent studies. Only two of the 81 analyses revealed significant results. We conclude the paper by discussing theoretical assumptions underlying the FAA and suggest a list of guidelines for improving and expanding the EEG data analysis in future FAA studies.

## Introduction

Electrophysiological studies on frontal alpha asymmetry (FAA) in depressive disorders (DD) have over 40 years of history, with first reports presented in 1979 by R. Davidson and colleagues (1979). Since then, many studies have reported relatively higher alpha band power in the left vs right frontal channels (left-sided FAA) in subjects suffering from DD compared to healthy individuals (Allen et al., 2004; Davidson, 1984, 2004; Kemp et al., 2010; Schaffer et al., 1983). FAA index, calculated by subtracting the left-side alpha power from the respective right-side channel, is one of the most common electrophysiological indicator of DD in the current literature (de Aguiar Neto & Rosa, 2019). However, multiple studies failed to replicate the relationship between FAA and DD (Allen et al., 2004; Carvalho et al., 2011; Deldin & Chiu, 2005; Gold et al., 2013; A. Kaiser et al., 2018; Kentgen et al., 2000; Knott et al., 2001; Mathersul et al., 2008; Szumska et al., 2020; Vuga et al., 2006) and conclusions of meta-analyses remain sceptical (Thibodeau et al., 2006; van der Vinne et al., 2017). In this light statements about FAA being a biomarker of depression (Baskaran et al., 2012; Iosifescu et al., 2009) seem to be too far-fetched.

It is not clear what the causes of above mentioned inconsistency in the literature are, but methodological issues are mentioned as one potential problem in a recent review by Kaiser et al. (2018). Although the authors focus mostly on the need to control for age, sex, handedness, medication and comorbidity, there are other important methodological problems worth considering when trying to resolve the validity of FAA in DD. Although much attention in the FAA literature has been paid to the choice of EEG reference (see for example: Smith et al., 2017; or Stewart et al., 2014) other aspects of signal processing and analysis seem to be more neglected. Many EEG studies on FAA use and report FAA index calculated only for a few channel pairs (e.g. one or two pairs were used in 12 out of 17 studies [70.6%] included in the meta-analysis by van der Vinne et al., 2017). In combination with the fact that topographical maps of effects are rarely presented (4/17 studies [23.5%] in van der Vinne et al., 2017) this significantly reduces the reliability and interpretability of the reported effects. The FAA effects are frequently assumed to reflect frontal sources of alpha oscillations but without a topographical map to support this claim it is difficult to conclude whether such interpretation is correct. For example alpha asymmetry at frontal channels may, in principle, arise due to asymmetrical projection from other, non-frontal, sources. This could be identified in the topography, but not at the single channel pair’s level. Without a topographical map it is also more difficult to assess the physiological reliability of the reported effect - significant effect on one channel pair without similar effects on surrounding channels calls for sceptical consideration (van Ede & Maris, 2016). Therefore, it might be better to perform the analysis on multiple frontal channel-pairs with relevant correction for multiple comparisons (for example with the very popular cluster-based permutation approach, Maris & Oostenveld, 2007). This is unfortunately rarely done in FAA studies on depressive disorders (0/17 studies in van der Vinne et al., 2017).

However, performing the analysis only at the channel level, when the research question pertains to the neural source of the effects, can also lead to misinterpretations. Even if topographies are shown, they can be inconclusive with respect to the underlying neural source. For this reason it might be useful to perform source localization and continue the analyses in the source space. Given the assumption of frontal alpha sources of FAA presented in the literature, source level analysis would be appropriate. Regrettably, most FAA studies do not perform source localization, although there are notable exceptions (for example: Lubar et al., 2003; Smith et al., 2018).

Incompatible results in FAA literature, summarized briefly above, suggest that the FAA relationship with DD is sensitive to the choice of signal preprocessing and analysis steps. In such a case applying multiverse analysis (Steegen et al., 2016), that is presenting results of multiple justifiable analysis paths, is a valuable tool to test the robustness of the studied effects. Multiverse analysis seems to be especially well suited for neuroscience research, given the multitude of preprocessing and data analysis choices that result in a complex “garden of forking paths” (Gelman & Loken, 2014). As most neuroscience studies test only one analysis variant it is difficult to assess the robustness of any individual effect, and it seems that at least some neuroscientific findings are sensitive to the choice of signal analysis steps (Cohen, 2015; Cohen & Gulbinaite, 2014; see also Botvinik-Nezer et al., 2019).

The purpose of this article is to thoroughly test the robustness and credibility of FAA as a marker of depressive disorders and address the limitations of FAA research methodology by performing a multiverse analysis of data coming from three independent studies. We performed 81 analyses in total differing in: a) the signal space used (channel space: average reference - AVG, current source density - CSD; source space: DICS beamforming); b) subselection of the signal space (channel pairs vs all frontal pairs with correction); c) statistical contrast used (group contrasts vs linear regression). Finally we propose additional guidelines to improve quality and reliability of the data analysis in FAA research.

## Methods

### Participants

In three independent studies we recruited 233 medication free participants in total - all without neurological disorders, drug or alcohol abuse, head injuries or stroke. Twelve subjects were excluded from further analyses due to excessive artifacts in the EEG signal (Study I: 0; Study II: 8; Study III: 4). Additional three subjects were excluded from Study I because of not fulfilling the analysis criteria. A total of 218 participants were included in the reported analyses: Study I, N = 51; Study II, N = 76; Study III, N = 91; for descriptives statistics summarizing each study see Table 1 and Figure 1.

**Figure 1.**
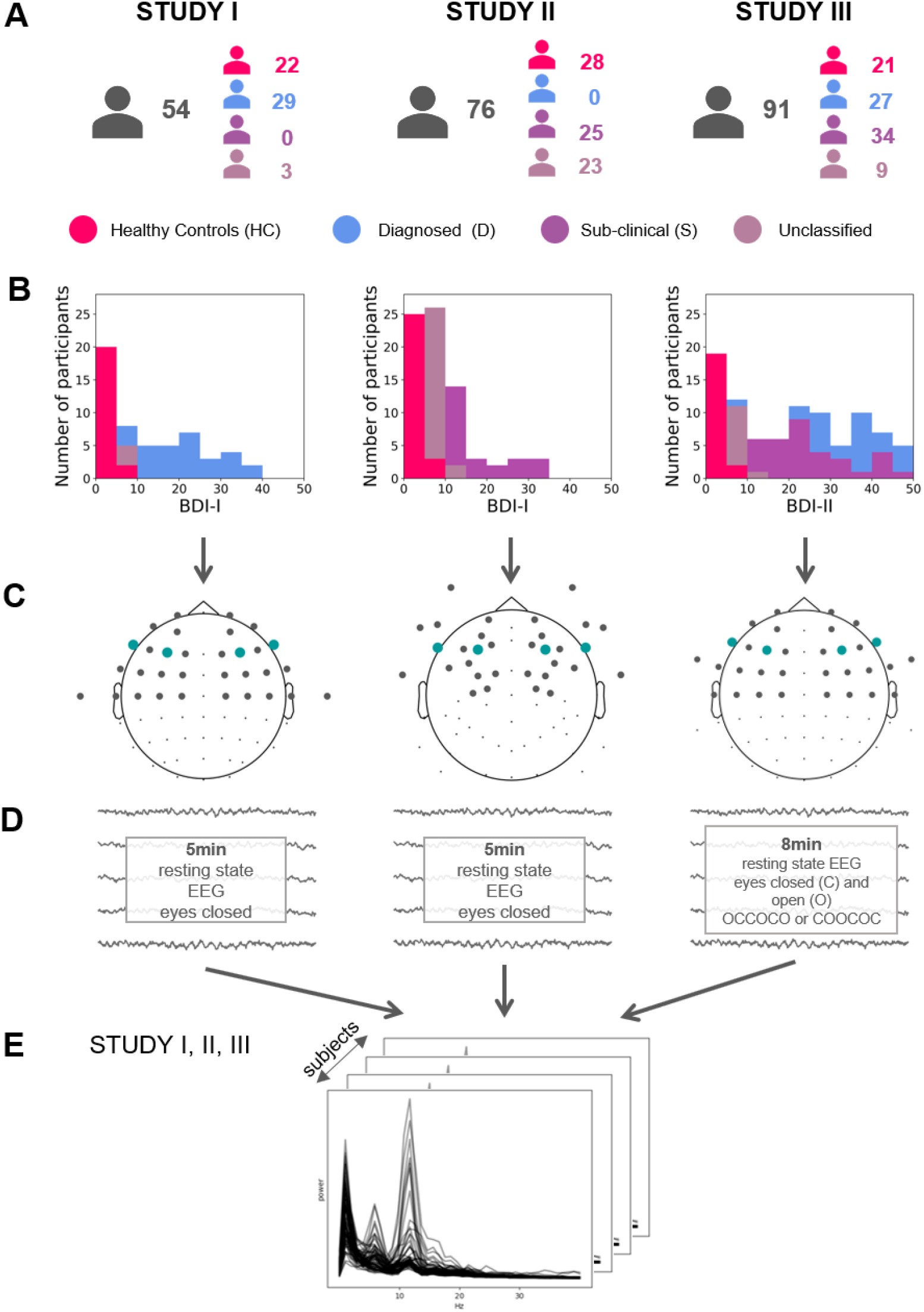
Diagram describing the three studies included in this article (Study I, II, III). (A) Number of participants for each study and group (see Table 1 for details). (B) Stacked histograms showing the distribution of BDI scores in each study and each group. (C) Channel montage. Frontal channels used in cluster based analyses are marked with gray dots. Channels used in channel-pairs analysis are marked with teal dots (F3 - F4, F7 - F8 and corresponding channels in the EGI montage). (D) Rest period length and scheme. (E) Channel-level spectra for each participant (see details in *Signal analysis* section).

**Table 1.** Descriptive statistics for each study presented in the article (*N - number of participants, M - mean score, SD - standard deviation, BDI-I and BDI-II - Beck Depression Inventory I and II*)

| Study I (N=51) |  |  |  |  |
| --- | --- | --- | --- | --- |
|  | Diagnosed | Healthy Controls, BDI-I <= 5 |  |  |
| N | 29 | 22 |  |  |
| Age | M = 27.66, SD = 7.13 | M = 23.68, SD = 2.83 |  |  |
| Gender | 9 male, 20 female | 11 male, 11 female |  |  |
| BDI-I score | M = 20.93, SD = 8.21 | M = 2.00, SD = 1.48 |  |  |
| Study II (N=76) |  |  |  |  |
| Undiagnosed (N=76) |  |  |  |  |
|  | Healthy Controls, BDI-I <= 5 | Sub-clinical, BDI-I >= 10 | Unclassified, 5 < BDI-I < 10 |  |
| N | 28 | 25 | 23 |  |
| Age | M = 25.32, SD = 6.46 | M = 24.44, SD = 5.08 | M = 25.22, SD = 6.78 |  |
| Gender | 8 male, 20 female | 4 male, 21 female | 9 male, 14 female |  |
| BDI-I score | M = 2.29, SD = 1.72 | M = 17.56, SD = 8.13 | M = 7.91, SD = 1.16 |  |
| Study III (N=91) |  |  |  |  |
|  | Diagnosed | Undiagnosed (N=64) |  |  |
|  |  | Healthy Controls, BDI-II <= 5 | Sub-clinical, BDI-II >= 10 | Unclassified, 5 < BDI-II < 10 |
| N | 27 | 21 | 34 | 9 |
| Age | M = 27.19, SD = 7.23 | M = 24.29, SD = 4.99 | M = 25.06, SD = 6.58 | M = 26.78, SD = 8.74 |
| Gender | 6 male, 21 female | 7 male, 14 female | 10 male, 24 female | 2 male, 7 female |
| BDI-II score | M = 34.26, SD = 9.18 | M = 2.24, SD = 1.70 | M = 24.06, SD = 10.08 | M = 6.78, SD = 0.97 |

Participants in Study I (only healthy controls, HC), II and III were recruited from the general population via advertisements in the local media or internal announcements for students at the University of Social Sciences and Humanities in Warsaw. In Study I where we had patients with diagnosis, they were recruited at the Psychiatry Clinic of the Department of Psychiatry, Medical University of Warsaw.

Each participant completed the Beck Depression Inventory (BDI) to determine current level of mood disorder: we used BDI version I (A. T. Beck et al., 1961) in Study I and II; and BDI version II (A. Beck et al., 1996) in Study III. Patients in Study I and III were diagnosed with mild depressive disorder (mild DD) (F32.0) according to ICD-10 classification criteria after a structured clinical interview using the MINI - mini-international neuropsychiatric interview (Sheehan et al., 1998). Local ethics committees (study I - from the Medical University of Warsaw, studies II and III - from the University of Social Sciences and Humanities) approved studies’ protocols and all participants signed consent forms.

### Data acquisition

The summary of all studies is presented in Figure 1. The equipment specifications and sessions details are provided below:

*Study I* - EEG signal was recorded with 64 channels (Ag/AgCl electrodes) arranged in the 10-5 system in a WaveGuard EEG Cap (Advanced Neuro Technology, ANT) at a sampling rate of 512 Hz. Impedance was kept below 10 kΩ. EEG signal was recorded during a five-minute session with eyes closed.
*Study II* - EEG signal was recorded with 64-Channel EGI HydroCel Geodesic Sensor Net, NetStation software and an EGI Electrical Geodesic EEG System 300 amplifier at a sampling rate of 200 Hz. Impedance was kept below 40 kΩ. EEG signal was recorded during a five-minute session with eyes closed.
*Study III* - EEG signal was recorded with 64-Channel (Ag/AgCl active electrodes) Brain Products ActiCap system and BrainVision software at a sampling rate of 1000 Hz and downsampled off-line to 250 Hz. Impedance was kept below 10 kΩ. EEG signal was recorded during an eight-minute session with alternating eyes open (O) and eyes closed (C) one minute segments. The ordering of the segments was either OCCOCO or COOCOC (chosen randomly for each participant).

### Data preprocessing

The preprocessing was performed with a custom-made EEGLAB-based MATLAB toolbox (eegDb: (Magnuski, 2020b)) and custom MATLAB scripts. Preprocessing steps were the same for all three studies. Continuous EEG signal was 1 Hz high pass filtered and divided into one-second consecutive segments. EEG recordings were visually inspected and segments containing strong or non-stereotypic artifacts were marked for rejection. These segments were ignored in all further preprocessing and analysis steps. Independent Component Analysis (ICA) was applied to remove artifacts related to eye blinks, eye movements, muscular and cardiac artifacts (Hipp & Siegel, 2013; McMenamin et al., 2010; Shackman et al., 2009). The average number of removed components in each study were as follows: M = 7.20, SD = 3.80 in *Study I*; M = 8.29, SD = 3.50 in *Study II*; M = 9.41, SD = 5.49 in *Study III*. Bad channels (*Study I:* M = 0.11, SD = 0.32; *Study II:* M = 0.72, SD = 1.04; *Study III:* M = 1.14, SD = 1.23) were not included in the ICA and were interpolated after cleaning the signal with ICA. The signal was re-referenced to common average (AVG) or current source density (CSD), depending on the type of analysis (see tables in *Results* section).

### Signal analysis

#### Channel-pair and cluster-based analyses

All channel and source level analyses were performed using mne-python (Gramfort et al., 2014) and custom code (Magnuski, 2020a, 2020c; Magnuski & Ruban, 2020; all available on github). Half of the analyses used current source density reference (CSD) and the other half used average reference (AVG; see Table *2* and Table *3* for a summary). For each dataset the continuous signal from eyes-closed rest period was used, starting 2 seconds after rest onset to avoid potential artifacts related to eyes closing. Power spectra were calculated using Welch method with 2-seconds long windows and a window step of 0.5 s. Welch windows overlapping with bad signal segments were removed and all remaining windows were averaged. This operation was performed for every channel and every subject giving rise to subjects by channels by frequencies matrix (Figure 1, panel E). Alpha asymmetry was calculated by first averaging spectral power in 8 - 13 Hz band, log-transforming and then for each left-right channel pair subtracting values obtained for left sites from those for right sites. We calculated alpha asymmetry as log(right) - log(left) because this is the most common approach.

#### Cluster-based analyses on standardized data

When the right-side alpha pattern is topographically different from the left-side alpha pattern we cannot expect left vs right subtraction to reliably uncover alpha asymmetry. To alleviate this problem we performed an additional analysis that does not rely on subtraction. In this approach all frontal channels were used including those at the midline. Moreover, the alpha frequency range (8 - 13 Hz) was not averaged, all frequency bins in this range were analyzed. Instead of right - left subtraction we standardized (z-scored) power in the selected channels by frequency space for each subject. Standardization should highlight asymmetry patterns that escape the classical left vs right comparison, while also being sensitive to effects that do not rely on asymmetry at all.

#### Source level analyses

Because channel-level projections can be highly variable depending on the source orientation we additionally perform analyses in the source space. We first digitized channel positions for a representative subject using photogrammetry. This step was performed because the default channel positions for many EEG caps assume a spherical head, which is not a realistic assumption for source localization. A hand-held video camera was used to record EEG cap placement on the head of the representative subject from multiple angles. The recorded video was processed with 3DF Zephyr (3DFlow 3DF Zephyr, Aerial Education version: Toldo et al., 2015) in order to obtain a 3D model of the subject’s head and EEG cap. Channels positions’ coordinates were extracted by manually placing control points on each channel in the 3d reconstruction. After coregistering the digitized channel positions with the fsaverage FreeSurfer head model (Dale et al., 1999), see next paragraph) we confirmed that the chosen subject’s head shape was very similar to the fsaverage head model.

We employed Boundary Element Method (BEM) for the forward problem. We first created a three-layer (inner skull, outer skull and outer skin) BEM model based on the FreeSurfer fsaverage template (Dale et al., 1999; Fischl et al., 1999). Next, the leadfield was constructed for a grid of 8196 equidistant source points covering the whole fsaverage cortical surface. Finally we used beamforming (Dynamic Imaging of Coherent Sources, DICS: Gross et al., 2001) to infer the source-level activity in alpha band.

The cross-spectral density matrices, necessary for DICS beamforming, were computed using Morlet wavelets (of length equal to seven cycles) on the continuous signal from the eyes-closed rest period starting from 2 seconds after rest onset. Bad signal segments were ignored, just like in the channel level analyses. To make the inverse solution more stable and noise-resistant we used a regularization parameter of 0.05 (van Vliet et al., 2018). Localized power maps were morphed to a symmetrical version of fsaverage brain (fsaverage_sym; Greve et al., 2013; Van Veen et al., 1997) to allow for left vs right comparisons. The asymmetry was computed in the same way as in the channel-pair and cluster-based analyses.

#### Statistical analysis

We performed a multiverse analysis consisting of 81 analyses differing in: a) the signal space used: channel space (average reference: 36 analyses, 44%, CSD reference: 36, 44%) or source space (DICS beamforming, 9, 12%); b) subselection of the signal space: channel pairs (36, 44%), all frontal pairs with correction (27, 34%) or all frontal channels without pairing with correction (18; 22%); c) statistical contrast used: group contrasts using independent t tests to compare FAA between groups; linear contrasts using linear regression to test the relationship between BDI scores and FAA. Group contrasts included: comparison between diagnosed and healthy controls (*DvsHC*) or sub-clinical and healthy controls (*SvsHC*); linear contrasts were performed either for all subjects together (*allReg*), only for diagnosed subjects (*DReg*) or only the non diagnosed subjects (*nonDReg*). These statistical contrasts are only used in the studies where they apply: for example contrasting healthy and diagnosed subjects (*DvsHC*) cannot be done for study II, where only healthy and sub-clinical participants are available. In the same way, comparing subclinical and healthy controls (*SvsHC*) is not possible in study I, where only healthy and diagnosed participants are available. The contrasts are performed for all analysis spaces: average reference, current source density and source level data. Additionally, for each statistical contrast and study we perform two data analysis approaches: a) classical comparison of selected two channel pairs; b) cluster-based permutation test on the whole frontal asymmetry space. The source space and analyses on standardized data employ only the cluster-based approach. Combining all these analytical pathways (*studies x analysis spaces x statistical contrasts x analysis approaches*) leads to 81 analyses: 54 come from the analyses in the channel space (*studies x channel spaces x contrasts*), 18 from *studies x channel spaces x contrasts* of the analyses on standardized data and 9 from *studies x contrasts* of the source space analyses. These analyses are summarized in Figure 2 and Tables 2–5.

**Figure 2.**
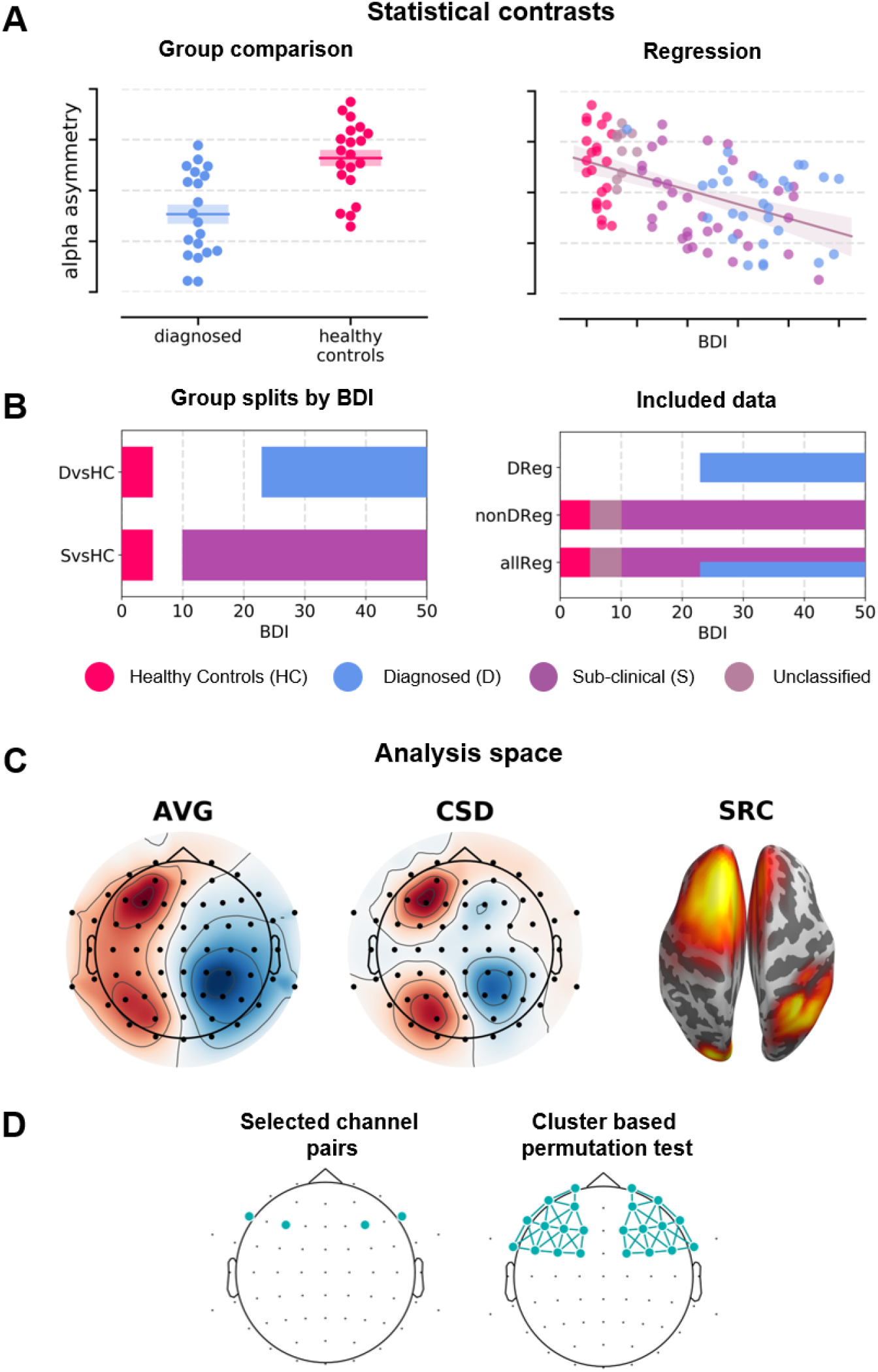
Analysis variants used (described in detail in *Statistical analysis* part) (A) Schematic depiction of given statistical contrast: group comparisons (left) vs regression (right). (B) Specification of each contrast against BDI scores. Left panel shows the range of BDI scores in each group in diagnosed vs healthy controls (*DvsHC*, first row) and diagnosed vs subclinical (*SvsHC*, second row) contrasts. Right panel shows the range of BDI scores for data included in each regression contrast: regression on diagnosed subjects (DReg) uses only subjects with clinical diagnosis, regression on non-diagnosed subjects (*NonDReg*) uses all subjects except those with clinical diagnosis; while regression on all subjects (*AllReg*) uses all subjects, hence all four colors are shown. The color legend for the subject groups is presented below these figures. (C) Analysis space: AVG - channel level, average reference; CSD - channel level, current source density; SRC - source level, DICS beamforming. (D) Schematic depiction of analysis method: selected channel pairs versus all frontal channels with cluster-based correction for multiple comparisons.

**Table 2.** Results for all channel-pair analyses. Each row represents two channel-pair results for a given contrast, study and space combination; uncorrected. Electrode placement for each study is shown in Figure 1C. (*N:* number of participants included in given contrast; *ES*: effect size; Cohen’s d for group comparison and Pearson’s r for regression; *CI*: bootstrap 95% confidence interval for the effect size).

| No. | CONTRAST | STUDY | SPACE | N | Selected channel-pairs without correction |  |  |  |  |  |  |  |
| --- | --- | --- | --- | --- | --- | --- | --- | --- | --- | --- | --- | --- |
|  |  |  |  |  | PAIR 1 (F3-F4) |  |  |  | PAIR 2 (F7 - F8) |  |  |  |
|  |  |  |  |  | t | p | ES | CI | t | p | ES | CI |
| 1 | DvsHC | I | avg | 29 vs 22 | -2.025 | 0.048 | -0.573 | [-1.135, -0.024] | -0.359 | 0.721 | -0.101 | [-0.644, 0.465] |
| 2 | DvsHC | I | csd | 29 vs 22 | 0.136 | 0.893 | 0.038 | [-0.550, 0.608] | 0.541 | 0.591 | 0.153 | [-0.415, 0.689] |
| 3 | DvsHC | III | avg | 27 vs 21 | 0.850 | 0.400 | 0.247 | [-0.316, 0.689] | -0.497 | 0.622 | -0.145 | [-0.760, 0.452] |
| 4 | DvsHC | III | csd | 27 vs 21 | 0.778 | 0.441 | 0.226 | [-0.307, 0.721] | -0.122 | 0.904 | -0.035 | [-0.590, 0.536] |
| 5 | SvsHC | II | avg | 23 vs 28 | 0.199 | 0.843 | 0.056 | [-0.502, 0.614] | -0.637 | 0.527 | -0.179 | [-0.780, 0.381] |
| 6 | SvsHC | II | csd | 23 vs 28 | 1.129 | 0.264 | 0.318 | [-0.221, 0.827] | -0.193 | 0.848 | -0.054 | [-0.611, 0.508] |
| 7 | SvsHC | III | avg | 33 vs 21 | -0.747 | 0.458 | -0.209 | [-0.640, 0.306] | -1.188 | 0.240 | -0.332 | [-0.804, 0.216] |
| 8 | SvsHC | III | csd | 33 vs 21 | -1.003 | 0.320 | -0.280 | [-0.730, 0.219] | -1.081 | 0.285 | -0.302 | [-0.798, 0.219] |
| 9 | allReg | III | avg | 91 | 0.545 | 0.587 | 0.058 | [-0.102, 0.263] | -0.781 | 0.437 | -0.083 | [-0.262, 0.112] |
| 10 | allReg | III | csd | 91 | 0.209 | 0.835 | 0.022 | [-0.171, 0.206] | 0.422 | 0.674 | 0.045 | [-0.161, 0.236] |
| 11 | DReg | I | avg | 29 | 0.980 | 0.336 | 0.185 | [-0.201, 0.544] | 0.320 | 0.751 | 0.061 | [-0.456, 0.534] |
| 12 | DReg | I | csd | 29 | -1.303 | 0.204 | -0.243 | [-0.552, 0.080] | -0.504 | 0.618 | -0.097 | [-0.503, 0.318] |
| 13 | DReg | III | avg | 27 | 0.278 | 0.784 | 0.055 | [-0.273, 0.438] | 0.055 | 0.957 | 0.011 | [-0.271, 0.244] |
| 14 | DReg | III | csd | 27 | 0.407 | 0.688 | 0.081 | [-0.260, 0.386] | 1.347 | 0.190 | 0.260 | [-0.076, 0.486] |
| 15 | nonDReg | II | avg | 76 | 0.223 | 0.824 | 0.026 | [-0.215, 0.238] | -0.760 | 0.450 | -0.088 | [-0.242, 0.078] |
| 16 | nonDReg | II | csd | 76 | 0.560 | 0.577 | 0.065 | [-0.132, 0.282] | -0.840 | 0.403 | -0.097 | [-0.275, 0.081] |
| 17 | nonDReg | III | avg | 64 | -0.796 | 0.429 | -0.101 | [-0.267, 0.112] | -1.599 | 0.115 | -0.199 | [-0.398, 0.039] |
| 18 | nonDReg | III | csd | 64 | -1.387 | 0.170 | -0.174 | [-0.399, 0.044] | -0.751 | 0.455 | -0.095 | [-0.342, 0.138] |

*DvsHC* - diagnosed and healthy controls; *SvsHC* - sub-clinical and healthy controls; *allReg* - linear regression for all subjects together; *DReg* - linear regression for only diagnosed subjects; *nonDReg* - linear regression for only the non diagnosed subjects; *avg* - average reference; *csd* - current source density

All the analyses that involve more than two selected channel pairs use cluster-based permutation tests (Maris & Oostenveld, 2007) to correct for multiple comparisons. Cluster based permutation test is a nonparametric multiple comparison correction where the hypothesis of difference between conditions is evaluated at the level of multidimensional clusters. Clusters are formed by performing a chosen statistical test in the n-dimensional search space (channels or channels by frequencies in most of the analyses reported here) and grouping adjacent points where the test statistic passed some predefined threshold (typically an alpha level of 0.05). Each obtained cluster is then summarized by summing the statistics of all its members - that is all adjacent points forming the cluster. These cluster summaries (cluster statistics) are then compared to a permutation null distribution of the maximum cluster statistic to obtain a p value. The null distribution is approximated by a Monte-Carlo method where in each draw condition labels are permuted between subjects (in this study: diagnosis status or BDI scores) and the statistical tests and clusters are computed in the same manner as for non-permuted data. As a result each Monte-Carlo draw produces cluster statistics from which the highest positive value and the lowest negative value are saved. These values, when aggregated from all Monte Carlo draws, constitute the null distribution for positive and negative effects to which cluster statistics from the actual analysis are compared.

For cluster-based analyses on standardized data, because they are sensitive to effects that do not have to be asymmetrical, significant test results were followed up with tests for asymmetry of the effects. For each cluster with p value below 0.05 a Chi-square test for two proportions was conducted comparing the proportion of cluster points on the left and right side of the head. Significant outcome of the test suggests that the cluster is asymmetrical.

Throughout all the analyses, including cluster-based permutation tests, we use an alpha level of 0.05. The same alpha level is used for cluster entry threshold in cluster-based tests. Results for single channel pairs, reported in Table 2, include also effect size (Cohen’s d for group comparisons, Pearson’s r for regression) and its 95% confidence interval calculated using bias-corrected accelerated bootstrap (Tibshirani & Efron, 1993; Ho et al., 2019).

## Results

### Channel-pair analyses

We reported results of all the channel-pair analyses in Table *2*. Only 1 out of 54 analyses gave significant results. This is expected by chance (p = 0.937, binomial test for a probability of significant result greater than 5%). Specifically, the significant result was found only for the F3 - F4 channel pair in the *DvsHC* contrast in study I with AVG reference. Alpha asymmetry was lower for the diagnosed group than for healthy controls, t = −2.025, p = 0.048 (Figure 3). To examine this result in a wider context, we compare it to the outcome of related analyses that differ in a single parameter. Applying CSD instead of average reference to the same channel pair did not reveal a significant result (same study, same contrast, F3 - F4 pair, CSD reference, t = 0.136, p = 0.893). The results were also not significant for the other channel pair, irrespective of reference (same study, same contrast, F7 - F8 pair: average reference, t = −0.359, p = 0.721; CSD reference, t = 0.541, p = 0.591). Performing the same analysis on data from study III also did not give rise to a significant outcome (same contrast, F3 - F4 pair, AVG reference, t = 0.85, p = 0.4; for other results see Table 2). For the study II the *DvsHC* contrast was not available, but conceptually closest contrast - Subclinical vs Healthy Controls (*SvsHC*) - was found insignificant for both channel pairs using either AVG or CSD references (see Table 2 for all results). However, as we argued in the introduction, single channel analyses are not a particularly good approach to FAA. They may be sensitive to small changes in the topography pattern and do not provide any information about the source or physiological plausibility of the effect.

**Figure 3.**
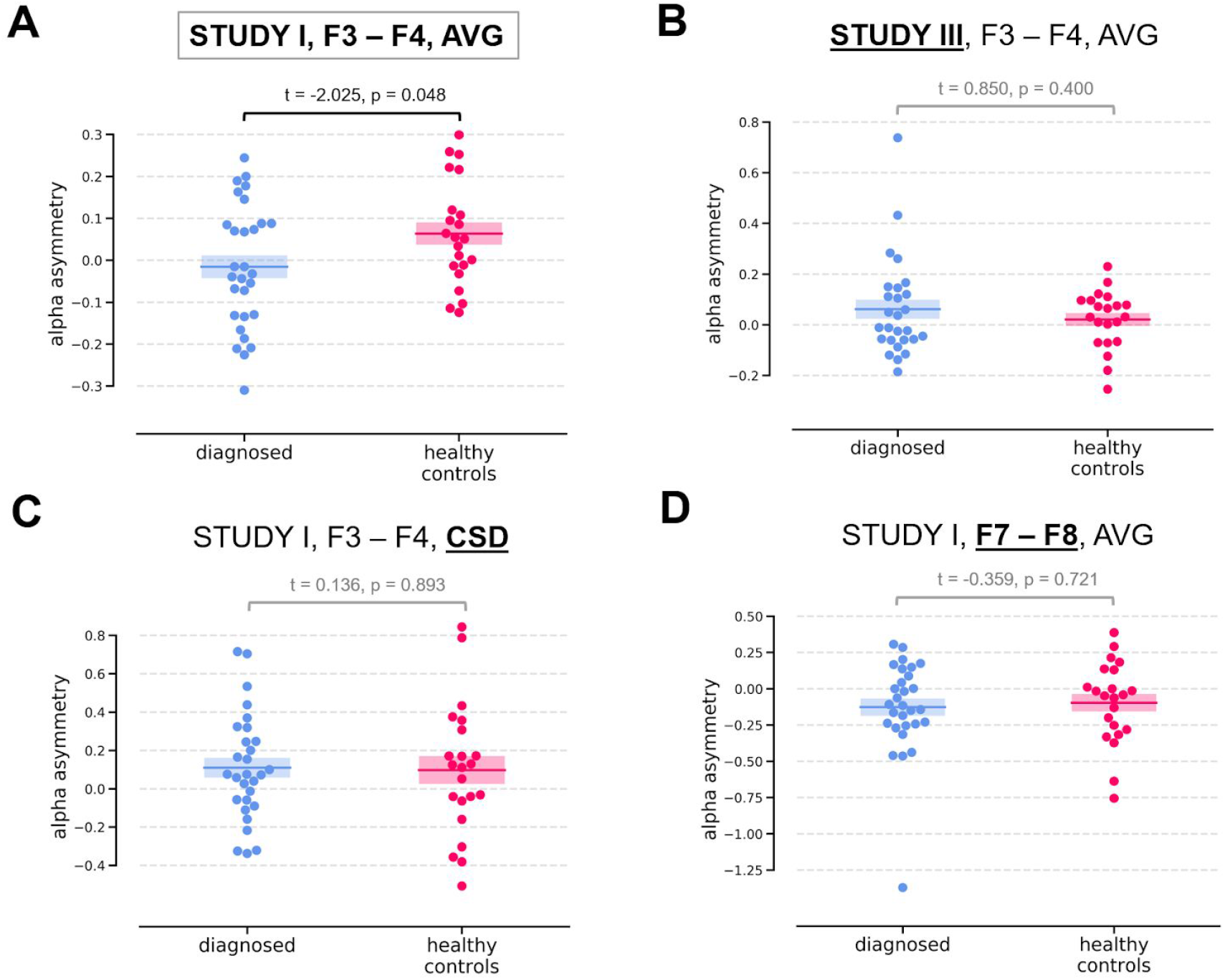
Selected results for channel-pairs level analyses,depressed versus healthy controls (*DvsHC*) contrast. Panel A (framed) shows the only significant channel pair result: study I, F3 - F4 channel pair, average reference. The remaining panels (B, C, D) show other channel pair analysis variants: specifically those that differ by exactly one parameter (underlined text) from the result presented in the panel A. Horizontal lines represent averages for each group, shaded areas show standard error of the mean. Results for study II are not presented in the figure, as *DvsHC* contrast was not available for this study.

### Cluster-based analyses

The next set of analyses consisted of cluster-based analyses on all frontal channel pairs. This approach gives a better view of the whole frontal alpha asymmetry space (correcting for multiple comparisons) especially when coupled with presentation of the effects’ topographies. Nonetheless we did not observe any significant effect in these analyses: either contrasting depressed (diagnosed or subclinical) vs healthy controls nor looking for a linear relationship between FAA and BDI (Table 3, Figure 4). Although single channels sometimes passed the significance threshold (see *n significant points* column in Table 3) these effects were not significant at the cluster level. In other words the clusters formed by these channels were not convincingly stronger from clusters observed under the null hypothesis (that is during the permutations).

**Figure 4.**
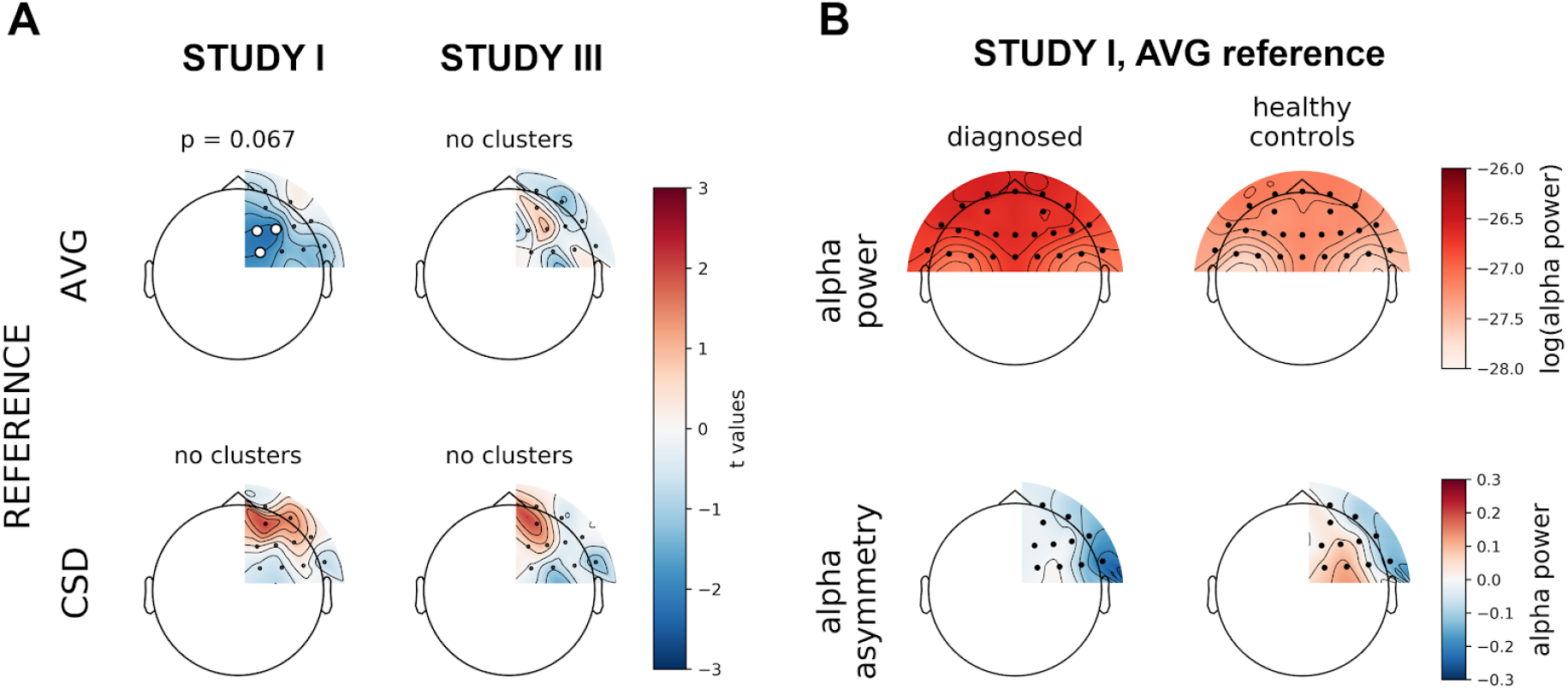
Selected results of cluster-based analyses. (A) topographies of *DvsHC* contrast effects in a reference by study matrix. More positive (red) t values indicate more right-sided (less left-sided) alpha asymmetry for diagnosed participants. More negative (blue) t values indicate more right-sided (less left-sided) FAA for healthy controls. Channels that are part of a cluster are marked with white dots. (B) Group-level averages for Study I, AVG from panel A. First row shows topography of average alpha power for both groups; second row shows topography of average alpha asymmetry for both groups. Results for study II are not presented, because this study did not include diagnosed subjects and *DvsHC* contrast was not possible.

**Table 3.** Results for cluster-based permutation test on frontal asymmetry space. Each row represents cluster-based results for a given contrast, study and space combination; (*N:* number of participants included in given contrast; *min t, max t*: lowest and highest t value in the search space, respectively; *n significant points:* total number of significant points in the search space before cluster-based correction; *n clusters:* number of clusters found in given analysis; *largest cluster size:* number of channels participating in the cluster; *largest cluster p:* p value for the largest cluster, *NA* means that no cluster was found in given analysis)

| No. | CONTRAST | STUDY | SPACE | N | Cluster-based analyses on frontal asymmetry space |  |  |  |  |  |
| --- | --- | --- | --- | --- | --- | --- | --- | --- | --- | --- |
|  |  |  |  |  | min t | max t | n significant points | n clusters | largest cluster size | largest cluster p |
| 1 | DvsHC | I | avg | 29 vs 22 | -2.094 | 0.022 | 3 | 1 | 3 | 0.067 |
| 2 | DvsHC | I | csd | 29 vs 22 | -0.720 | 1.964 | 0 | 0 | NA | NA |
| 3 | DvsHC | III | avg | 27 vs 21 | -1.159 | 0.850 | 0 | 0 | NA | NA |
| 4 | DvsHC | III | csd | 27 vs 21 | -1.257 | 1.758 | 0 | 0 | NA | NA |
| 5 | SvsHC | II | avg | 23 vs 28 | -2.571 | 0.199 | 1 | 1 | 1 | 0.137 |
| 6 | SvsHC | II | csd | 23 vs 28 | -0.827 | 1.229 | 0 | 0 | NA | NA |
| 7 | SvsHC | III | avg | 33 vs 21 | -1.318 | 0.930 | 0 | 0 | NA | NA |
| 8 | SvsHC | III | csd | 33 vs 21 | -2.294 | 1.877 | 1 | 1 | 1 | 0.214 |
| 9 | allReg | III | avg | 91 | -1.207 | 0.906 | 0 | 0 | NA | NA |
| 10 | allReg | III | csd | 91 | -1.095 | 2.210 | 1 | 1 | 1 | 0.262 |
| 11 | DReg | I | avg | 29 | -0.159 | 1.552 | 0 | 0 | NA | NA |
| 12 | DReg | I | csd | 29 | -1.303 | 0.487 | 0 | 0 | NA | NA |
| 13 | DReg | III | avg | 27 | -2.041 | 0.976 | 1 | 0 | NA | NA |
| 14 | DReg | III | csd | 27 | -1.223 | 1.662 | 0 | 0 | NA | NA |
| 15 | nonDReg | II | avg | 76 | -2.004 | 0.751 | 1 | 1 | 1 | 0.342 |
| 16 | nonDReg | II | csd | 76 | -0.840 | 1.215 | 0 | 0 | NA | NA |
| 17 | nonDReg | III | avg | 64 | -1.599 | 1.387 | 0 | 0 | NA | NA |
| 18 | nonDReg | III | csd | 64 | -1.634 | 1.799 | 0 | 0 | NA | NA |

*DvsHC* - diagnosed and healthy controls; *SvsHC* - sub-clinical and healthy controls; *allReg* - linear regression for all subjects together; *DReg* - linear regression for only diagnosed subjects; *nonDReg* - linear regression for only the non diagnosed subjects; *avg* - average reference; *csd* - current source density

The only result close to our significance threshold was *DvsHC* contrast in study I using average reference (cluster p = 0.067). This is the same analysis combination as the significant channel-pair result reported in the previous section and similarly all comparable cluster-based analyses do not replicate it: see Figure 4 for a comparison between this cluster-based effect and other analyses differing by one parameter.

### Cluster-based analyses on standardized data

All previous analyses assume that asymmetry can be detected by subtracting right and left homologous channels. Because some asymmetry effects may not match such strict left vs right pattern, we performed further analyses to alleviate this issue: in this set of analyses we relied on standardization of alpha power at frontal channels instead of right minus left subtraction. Additionally, to minimize the risk of averaging out an effect confined to a narrow frequency range, we also analysed all frequency bins in the 8 to 13 Hz range. This strategy is used in a set of 18 analyses - we reported their results in *Table 4*.

**Table 4.** Results for all cluster-based analyses on standardized data. Each row represents cluster-based results for given contrast, study and space; (*N:* number of participants included in given contrast). Results for cluster-based permutation test on frontal asymmetry space (*min t, max t*: lowest and highest t value in the search space, respectively; *n significant points:* total number of significant points in the search space before cluster-based correction; *n clusters:* number of clusters found in given analysis; *largest cluster size:* number of channels by frequency points participating in the cluster; *largest cluster p:* p value for the largest cluster, *NA* means that no cluster was found in given analysis).

| No. | CONTRAST | STUDY | SPACE | N | Cluster-based analyses on standardized data |  |  |  |  |  |
| --- | --- | --- | --- | --- | --- | --- | --- | --- | --- | --- |
|  |  |  |  |  | min t | max t | n significant points | n clusters | largest cluster size | largest cluster p |
| 1 | DvsHC | I | avg | 29 vs 22 | -1.433 | 1.425 | 0 | 0 | NA | NA |
| 2 | DvsHC | I | csd | 29 vs 22 | -2.223 | 1.941 | 0 | 2 | 2 | 0.687 |
| 3 | DvsHC | III | avg | 27 vs 21 | -2.284 | 2.215 | 0 | 3 | 2 | 0.573 |
| 4 | DvsHC | III | csd | 27 vs 21 | -2.527 | 3.469 | 0 | 3 | 15 | 0.140 |
| 5 | SvsHC | II | avg | 23 vs 28 | -1.647 | 2.186 | 0 | 1 | 1 | 0.742 |
| 6 | SvsHC | II | csd | 23 vs 28 | -1.919 | 1.817 | 0 | 0 | NA | NA |
| 7 | SvsHC | III | avg | 33 vs 21 | -2.166 | 1.688 | 0 | 1 | 2 | 0.601 |
| 8 | SvsHC | III | csd | 33 vs 21 | -2.166 | 2.707 | 0 | 5 | 2 | 0.775 |
| 9 | allReg | III | avg | 91 | -2.076 | 2.055 | 0 | 2 | 1 | 1.000 |
| 10 | allReg | III | csd | 91 | -2.310 | 2.115 | 0 | 3 | 4 | 1.000 |
| 11 | DReg | I | avg | 29 | -2.011 | 1.936 | 0 | 0 | NA | NA |
| 12 | DReg | I | csd | 29 | -2.721 | 2.093 | 0 | 5 | 3 | 1.000 |
| 13 | DReg | III | avg | 27 | -4.086 | 3.661 | 1 | 3 | 73 | 0.024 |
| 14 | DReg | III | csd | 27 | -3.452 | 3.400 | 0 | 6 | 21 | 0.143 |
| 15 | nonDReg | II | avg | 76 | -1.740 | 1.307 | 0 | 0 | NA | NA |
| 16 | nonDReg | II | csd | 76 | -1.797 | 1.450 | 0 | 0 | NA | NA |
| 17 | nonDReg | III | avg | 64 | -2.466 | 2.183 | 0 | 3 | 2 | 1.000 |
| 18 | nonDReg | III | csd | 64 | -1.822 | 2.857 | 0 | 1 | 3 | 1.000 |

*DvsHC* - diagnosed and healthy controls; *SvsHC* - sub-clinical and healthy controls; *allReg* - linear regression for all subjects together; *DReg* - linear regression for only diagnosed subjects; *nonDReg* - linear regression for only the non diagnosed subjects; *avg* - average reference; *csd* - current source density

Only one standardization analysis showed statistically significant result - this is expected by chance given our alpha level (p = 0.603, binomial test). Specifically, the significant result was found for linear relationship with BDI confined to diagnosed subjects (*DReg* contrast) in study III using average reference (cluster p = 0.024). This effect is potentially interesting - it shows a positive relationship between depression severity (BDI score, only the diagnosed subjects) and power in the higher alpha band (11 - 12.5 Hz) across many frontal channels (see Figure 5). Because it is accompanied with an inverse effect in a lower frequency (9 - 10 Hz) at similar channels, averaging across the whole 8 - 13 Hz frequency range could lead to both effects cancelling each other in the average. Such a pattern of inverse effects across frequencies could arise due to a frequency shift of the individual alpha peak or the narrowing of the peak with depression severity. However, the effect does not seem to represent FAA as its topography is symmetrical (*χ*^2^(1) = 0.045, p = 0.831) and was not replicated when using CSD reference in the same study (cluster p = 0.143) or performing the same analysis on study I data (AVG: no clusters found; CSD: cluster p = 1.0).

**Figure 5.**
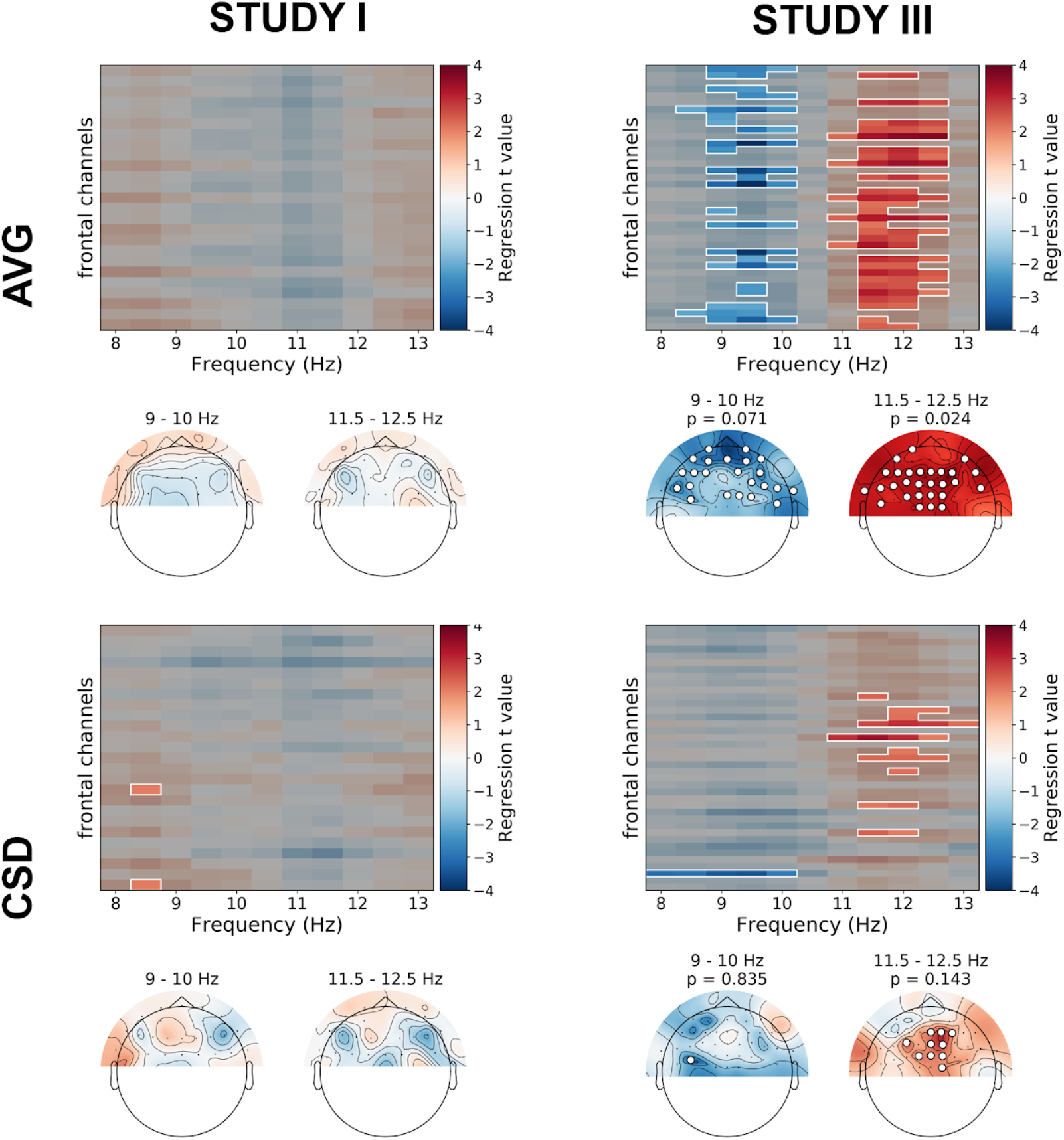
Selected results of cluster-based analyses on standardized data, DReg contrast (regression between FAA and BDI restricted to diagnosed subjects). Heatmaps in the upper part of each panel represent regression t-values for channel by frequency search space. More positive / negative t values, indicate higher / lower power with higher BDI. Clusters are indicated in the heatmaps with white outline. In each panel we present two topographies below the heatmap: showing average effect for 9 - 10 Hz and 11.5 - 12.5 Hz range respectively. Channels that are part of a cluster are marked with white dots in the topographical plots. Results for study II are not presented, because this study did not include diagnosed subjects so DReg contrast was not possible.

Although the conclusions that can be drawn from these standardization analyses are not in favor of FAA - DD relationship, they demonstrate the strength of the proposed approach in detecting effects that might be otherwise missed when averaging across the whole alpha range or when testing only the differences on corresponding right - left channel pairs.

### Source level analyses

Because observing the effect in the signal recorded from frontal channels does not guarantee that the source of this effect is frontal we conducted a second set of additional analyses using source localization with DICS beamforming. In these analyses the FAA was evaluated in the source space by subtracting power of the corresponding right and left hemisphere vertices.

Results of source level analyses are reported in Table 5 and Figure 6. None of the source space analyses turned out statistically significant - this is expected by chance given our alpha value (p=1., binomial test). In most of the analyses the pattern and sign of the t values points towards a more left-sided effect. For example in the linear regression for all subjects (*allReg* contrast): the negative t values suggest lower R - L differences in high than in low BDI participants, which means more left-sided alpha power with higher BDI. Although this pattern seems to be in line with FAA - DD literature, all of the source space effects are weak and do not survive the correction for multiple comparisons. However, it is important to remember that individual MRI scans were not available in the present study: their availability would lead to lower error in source reconstruction.

**Figure 6.**
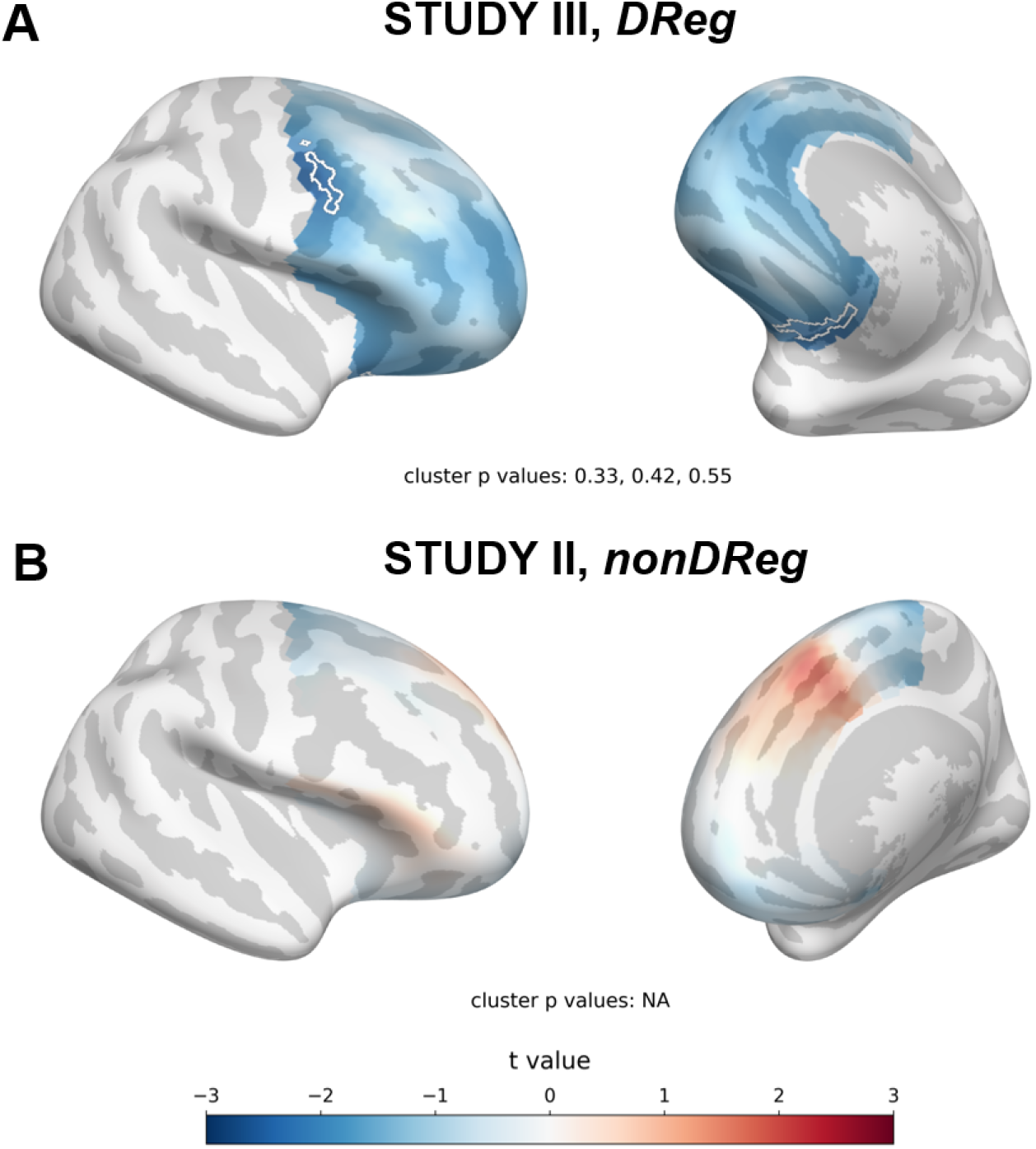
Selected results of source level analyses showing spatial t-value maps for regression analyses. Cluster limits are marked with white outlines, corresponding cluster p values are shown below each panel. Colorbar at the bottom presents color coding for the t values: more positive (red) / negative (blue) relationship between FAA and BDI. (A) Results for study III, *DReg contrast* - linear regression restricted to diagnosed subjects; no cluster with p < 0.05 is present. (B) Results for study II, *nonDReg* contrast - linear regression restricted to the non diagnosed subjects; no custer is present.

**Table 5.** Results for all source level analyses. Each row represents source level results for given contrast, study and space; (*N*: number of participants included in given contrast) Results for cluster-based permutation test on frontal asymmetry source space (*min t, max t*: lowest and highest t value in the search space, respectively; *n significant points:* total number of significant points in the search space before cluster-based correction; *n clusters:* number of clusters found in given analysis; *largest cluster p:* p value for the largest cluster, *NA* means that no cluster was found in given analysis).

| No. | CONTRAST | STUDY | N | Source level analysis |  |  |  |  |  |
| --- | --- | --- | --- | --- | --- | --- | --- | --- | --- |
|  |  |  |  | min t | max t | n significant points | n clusters | largest cluster size | largest cluster p |
| 1 | DvsHC | I | 29 vs 22 | -1.818 | 2.368 | 2 | 1 | 2 | 0.493 |
| 2 | DvsHC | III | 27 vs 21 | -2.541 | 1.167 | 28 | 4 | 11 | 0.287 |
| 3 | SvsHC | II | 23 vs 28 | -0.923 | 1.327 | 0 | 0 | NA | NA |
| 4 | SvsHC | III | 34 vs 21 | -1.762 | 1.195 | 0 | 0 | NA | NA |
| 5 | allReg | III | 92 | -2.232 | 0.290 | 19 | 2 | 20 | 0.353 |
| 6 | DReg | I | 29 | -2.292 | 1.150 | 54 | 2 | 22 | 0.335 |
| 7 | DReg | III | 27 | -2.624 | -0.375 | 33 | 3 | 18 | 0.331 |
| 8 | nonDReg | II | 76 | -1.686 | 1.956 | 0 | 0 | NA | NA |
| 9 | nonDReg | III | 65 | -1.725 | 1.003 | 0 | 0 | NA | NA |

*DvsHC* - diagnosed and healthy controls; *SvsHC* - sub-clinical and healthy controls; *allReg* - linear regression for all subjects together; *DReg* - linear regression for only diagnosed subjects; *nonDReg* - linear regression for only the non diagnosed subjects

## Discussion

We conducted a multiverse analysis of EEG data from three independent studies, with 218 participants and 81 analyses in total to test the robustness and credibility of the relationship between frontal alpha asymmetry and depressive mood. We performed 36 replicatory single channel pairs analyses and 18 corresponding cluster-based analyses. We have also conducted 27 additional analyses addressing some of the limitations of current FAA studies. Out of 81 performed analyses only two produced statistically significant results - a result expected by chance with 0.05 alpha level (binomial test, p = 0.917). Overall, the conducted analyses do not provide a basis to reject the null hypothesis of no relationship between resting state FAA and DD.

Our conclusion is similar to this formulated by other research groups (A. K. Kaiser et al., 2018; van der Vinne et al., 2017), stating that treating FAA as a biomarker of DD is not sufficiently empirically grounded. As our data shows, this skepticism is not limited to single channel pair analyses - improving on the limitations of methods commonly used in FAA literature does not change the pattern of results.

Despite this, FAA is one of the most common indicators of DD with a long history of successful studies - it might be difficult to believe that all previous FAA research represents Type I errors. Therefore it’s worth considering that the FAA - DD relationship exists but we fail to detect this effect here. First, the FAA effect size may be too small to be reliably observed with a small to moderate sample size. The average number of participants per study is 73 in our case, but some analyses contain around 30 participants per group, which grants sufficient power to detect mostly large effects. Although single analyses reported here can be deemed inconclusive the whole multiverse set of analyses is incompatible with presence of moderate to strong relationship between FAA and DD. To strengthen this point we show effect sizes and their confidence intervals for aggregated studies in *Figure 3 - figure supplement 1 and 2:* bayes factors presented there indicate that there is moderate evidence for the null hypothesis in 19 of 20 aggregated single-channel analyses. Although we cannot exclude a small to moderate FAA - DD relationship - if this effect was in this range then most published studies would have been underpowered to detect it. This line of thought is also supported in the meta-analysis by van der Vinne et al. (2017) - which shows that studies with larger samples were less likely to report high effect sizes. For example the largest EEG FAA study on a sample of 1008 DD patients and 336 controls didn’t confirm the diagnostic value of FAA in DD (Arns et al., 2016). Such pattern of results suggest publication bias or that the FAA - DD effect, if it exists, is detectable only in highly selected samples and is of small magnitude on the population level.

Smith et al. (2017) previously suggested that the relationship between FAA and DD is stronger when the participant is given some emotion-related task, as opposed to resting condition, where the task is unspecified. Although this is possible, we wanted to stay true to the design of most FAA - DD studies, which measure EEG during rest. An interesting approach for future studies would be to compare rest blocks separated by an emotional task (see for example: Beeney et al., 2014).

Another explanation of our null results might be due to the fact that our depressed groups consisted of participants diagnosed with mild depressive disorder (study I and III) or classified as subclinical on the basis of BDI test score (study II). DD symptoms for such patients are weaker and it could be argued that therefore the FAA - DD relationship may be more difficult to observe. On the other hand, clinical participants in studies I and III and subclinical participants in studies II and III manifested a wide range of DD symptoms indicated by BDI scores (see Figure 1 for BDI histograms), but regression analysis (*DReg* and *nonDReg* contrasts) did not reveal any relationship between FAA and BDI score. This means that participants with stronger DD symptoms were not better characterised by a specific FAA pattern.

A challenge for future FAA studies would be to move beyond a “marker-only” approach, describe the theoretical assumptions behind FAA in more detail and let these assumptions dictate an adequate analytical approach. For example, the assumption that FAA is a phenomenon with a source in the frontal cortex is impossible to address using only a few frontal channel pairs. Using source localization (like DICS Beamforming used here) or source separation (for example spatial filtering with generalized eigendecomposition / common spatial pattern, (Koles et al., 1990; Parra & Sajda, 2003; Tomé, 2006); or SPACE decomposition, (R. van der Meij et al., 2015; Roemer van der Meij et al., 2016) should be preferred when looking for the answer to this question. This point is important because frontal alpha sources are rarely measured reliably with EEG at the channel level: strong occipital and parietal alpha sources dominate alpha power recorded at frontal channels. As a result measuring FAA at the channel level could lead to poor signal to noise ratio and consequently to small effect size and low probability to observe a true FAA - DD relationship. On the other hand, if FAA does not originate in the frontal regions, it should be measured and interpreted differently. Such scenario is not unlikely because frontal alpha sources are generally difficult to detect with EEG/MEG. For example (Roemer van der Meij et al., 2016), using an advanced source separation method, found that 86.6% of the alpha components detected across subjects were occipito-parietal and only 1% (4 / 380) were frontal. If frontal alpha sources are difficult to detect using a source separation method designed to capture oscillatory sources, then it is likely that frontal alpha sources are rarely observed at the channel level at all.

If FAA is not of frontal origin then where could it come from? This is an open empirical question and we can only offer speculation here. Jiang et al. (2016) have shown that the power of posterior alpha oscillations is reduced in depressed individuals and that this reduction strongly correlates with depression severity. Assuming that frontal projections from occipital or parietal sources will not be perfectly symmetrical - a difference metric like FAA may be sensitive to posterior alpha power. Another possibility is that FAA originates from asymmetry at the source level: Smith et al. (2018) demonstrated that channel-level FAA is related to source level asymmetry in frontal motor regions. However, the authors didn’t look for correlations with FAA in the full source space, but restricted their analyses to the R - L source level asymmetry and in consequence their analysis is insensitive to symmetrical sources of FAA. Our data and analyses can’t be conclusive in this regard because without a strong and reliable FAA - DD effects it is difficult to look for it’s source level correlates. Tackling all the mentioned issues would require to systemize and unify the FAA methodology for the benefit of future studies. So far, Kaiser et al. (2018) proposed guidelines for methodology regarding subjects selection and controlling for confounding variables. Smith et al. (2017) also suggested possible improvements in experimental procedures and EEG signal preprocessing. We believe there is still room for improvement in the signal analysis standards of FAA studies. Below we summarize our arguments and propose additional guidelines for EEG data analysis in FAA research:

- Always show the topography of the effects. Lack of topographical plots hinders interpretation both in terms of potential neural origin of the effect and its physiological reliability. It is a good idea to also add topographical plots of group averages: both for alpha power and alpha asymmetry (see Figure 4 B). Such visualisations can clarify the studied effect: when FAA is calculated as a R - L difference and is compared between groups, reasoning about difference between differences ((R - L) - (R - L)) can be unnecessarily complex.
- Conduct analysis on all frontal electrodes (or even all available electrodes) with correction for multiple comparisons. We recommend using the cluster based permutation test (Maris & Oostenveld, 2007), as it is versatile and implemented in multiple software packages: mne-python (Gramfort et al., 2013, 2014) and fieldtrip (Oostenveld et al., 2011) for example, but the fieldtrip implementation is available also through EEGLAB (Delorme & Makeig, 2004) and brainstorm (Tadel et al., 2011).
- Try not to restrict the analysis to left minus right subtraction on averaged frequencies. As we show in additional analyses I - avoiding subtraction and frequency averaging can uncover interesting effects that could otherwise be missed. Extending the search space to frequencies is straightforward when using the cluster based permutation test.
- Perform analysis in the source-space if possible. Source localization allows to estimate the source of the signal more reliably and obtain a better signal to noise ratio. However, to minimize source localization error individual MRI scans are required. Other methods focusing on source-separation like ICA, GED or SPACE allow to disentangle signal contributions from independent sources and increase signal to noise ratio. CSD was also proposed in this context in FAA literature before, but although it can mitigate some of the issues arising from volume conduction, it does not provide source localization.
- Do not restrict the analysis to group contrasts if linear predictors are available. Using linear regression allows to take covariates into account and test hypotheses in a more detailed manner.

## Acknowledgements

The work was supported by the grants from the Ministry of Science and Higher Education: “Diamond Grant” (DI2013012943), Iuventus Plus grant (0045/IP3/2011/71) and N10601731/1344 grant. We thank Paweł Wroński for data collection in Study II; Paweł Holas and Dorota Wołyńczyk - Gmaj for psychiatric support in studies I and III.

We acknowledge the support of COVID-19 pandemic in keeping us locked in homes with nothing more interesting to do but writing this manuscript.

## Supplemental Figures

**Figure 3 - supplements 1 and 2:**

To strengthen our message of “no relationship” between FAA and DD we perform additional analyses for single channel pairs where we aggregate studies including identical contrasts, calculate the resulting effect size (Cohen’s d for group comparisons and Pearson’s r for linear relationships), estimate confidence intervals for the effect size using bias-corrected bootstrapping (Tibshirani & Efron, 1993; Ho et al., 2019) and calculate bayes factor for the null hypothesis (BF01; Rouder et al., 2009) using Pingoin python package using default settings (Vallat, R., 2018). In all group contrasts the confidence intervals exclude effect sizes of 0.6 d and higher (and −0.6 and lower) and BF01 indicate moderate evidence for the null hypothesis. In all linear relationships except one (DReg, AVG reference, channel pair F3 - F4) the confidence intervals exclude effect sizes of 0.35 r and higher (and −0.35 and lower) and BF01 indicate moderate evidence for the null hypothesis.

**Figure 3 - figure supplement 1:**
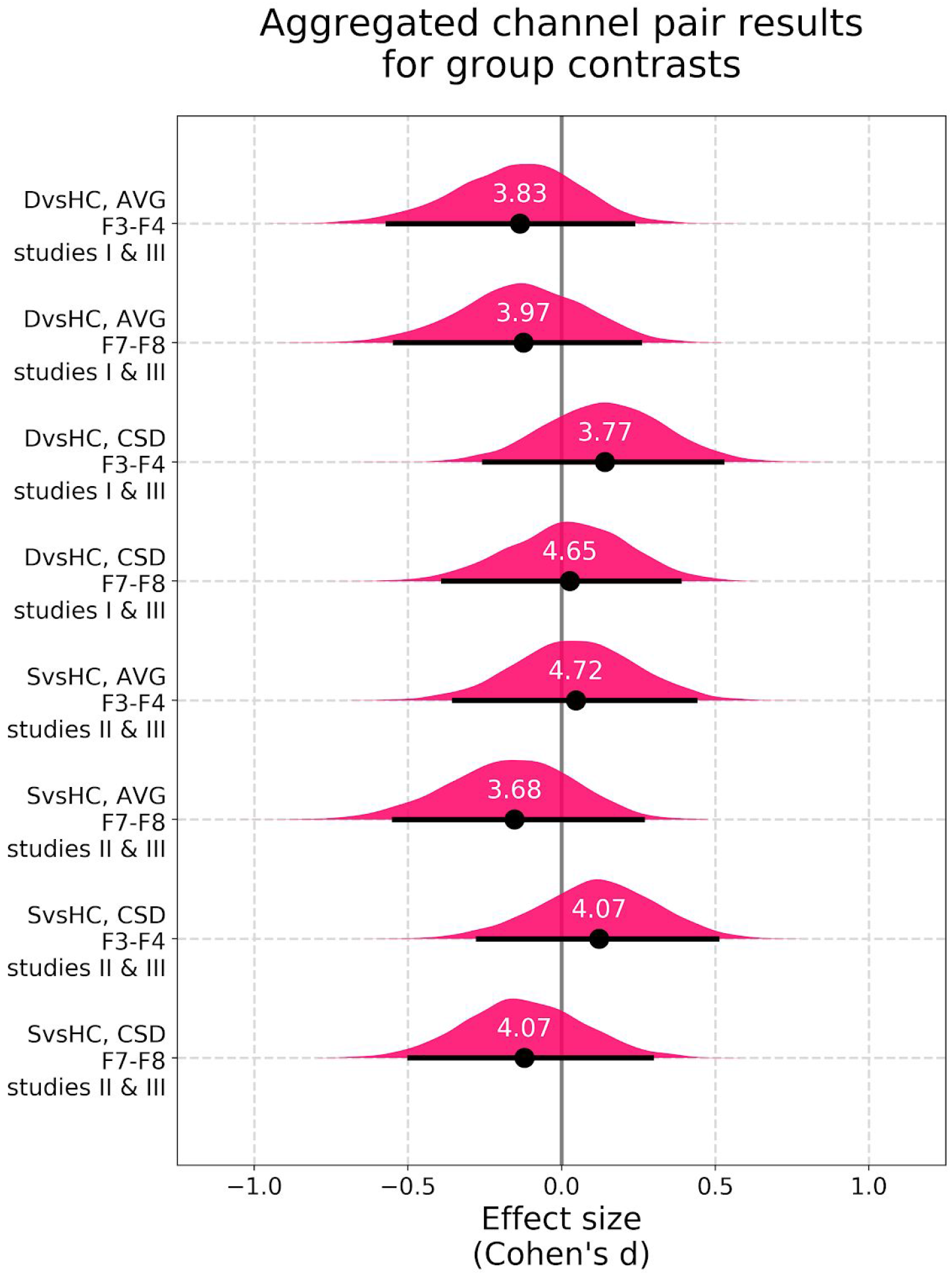
Results for aggregated analyses where studies including identical group contrasts are combined. Each row corresponds to one analysis on a single channel pair. The contrasts, studies and channel pairs are labeled on the y axis. The contrasts are abbreviated as in the main text: DvsHC: depressed individuals vs healthy controls; SvsHC: subclinical individuals (high BDI but no clinical diagnosis) vs healthy controls. The black dots correspond to observed effect sizes in Cohen’s d, while the black lines indicate 95% confidence intervals for the effect size estimated using bias-corrected accelerated bootstrapping. The magenta shapes represent distributions from the bootstrap and the white numbers printed on the magenta shapes are bayes factors for the null hypothesis (BF01). BF01 of 4 indicates that the data are four times more likely under the null than the alternative hypothesis. BF01 between 3 and 10 are considered moderate evidence for the null hypothesis.

**Figure 3 - figure supplement 2:**
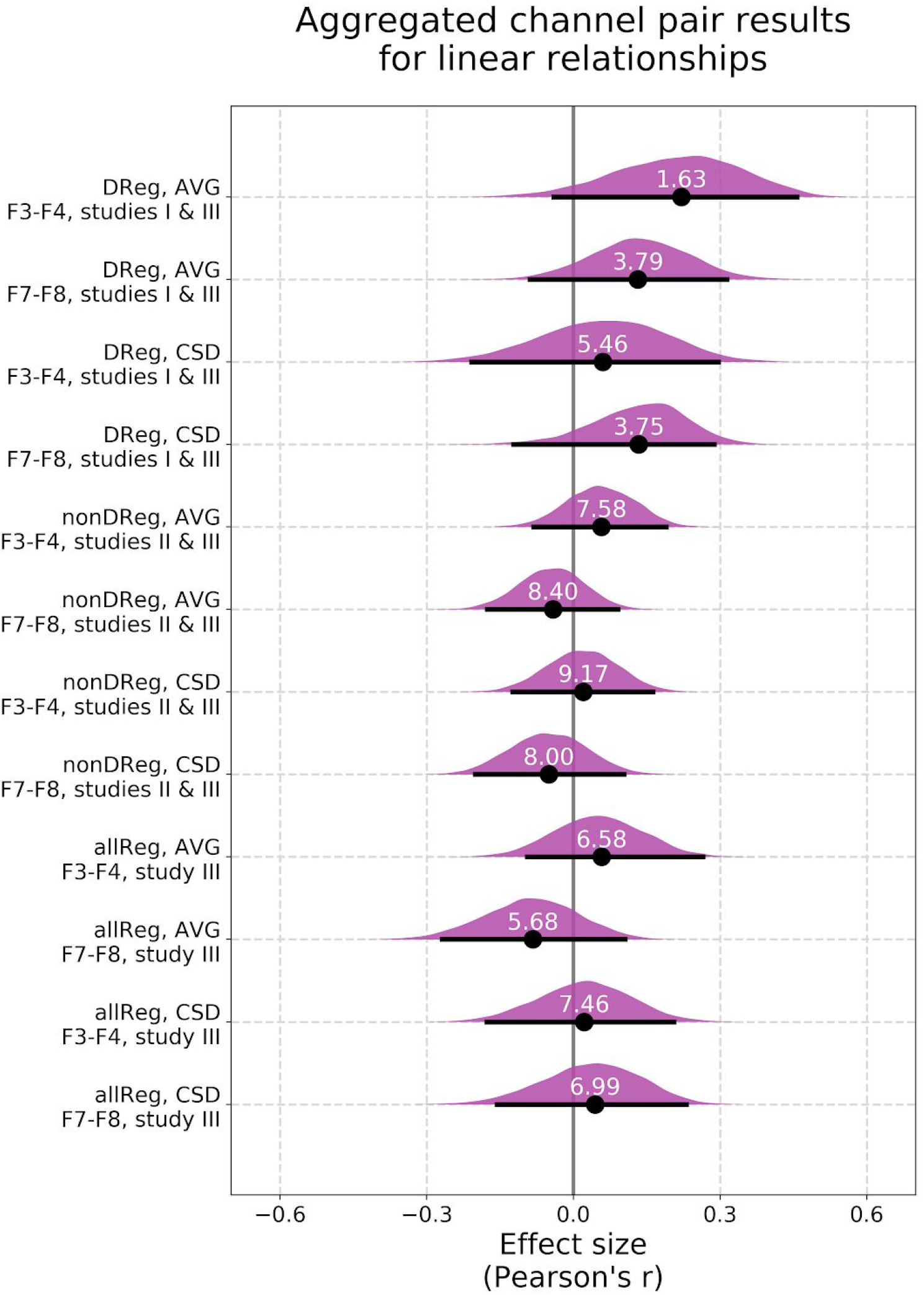
Results for aggregated analyses where studies including identical linear contrasts are combined. Each row corresponds to one analysis on a single channel pair. The contrasts, studies and channel pairs are labeled on the y axis. The contrasts are abbreviated as in the main text: DReg: linear relationship between FAA and BDI restricted to depressed individuals; *nonDReg*: linear relationship between FAA and BDI restricted to individuals without diagnosed depression; allReg: linear relationship between FAA and BDI for all participants. The black dots correspond to observed effect sizes in Pearson’s r, while the black lines indicate 95% confidence intervals for the effect size estimated using bias-corrected accelerated bootstrapping. The lavender shapes represent distributions from the bootstrap and the white numbers printed on the lavender shapes are bayes factors for the null hypothesis (BF01). BF01 of 4 indicates that the data are four times more likely under the null than the alternative hypothesis. BF01 between 3 and 10 are considered moderate evidence for the null hypothesis.

**Figure 4 - supplements 1 - 5: All results of cluster-based analyses.**

**Figure 4 - figure supplement 1:**
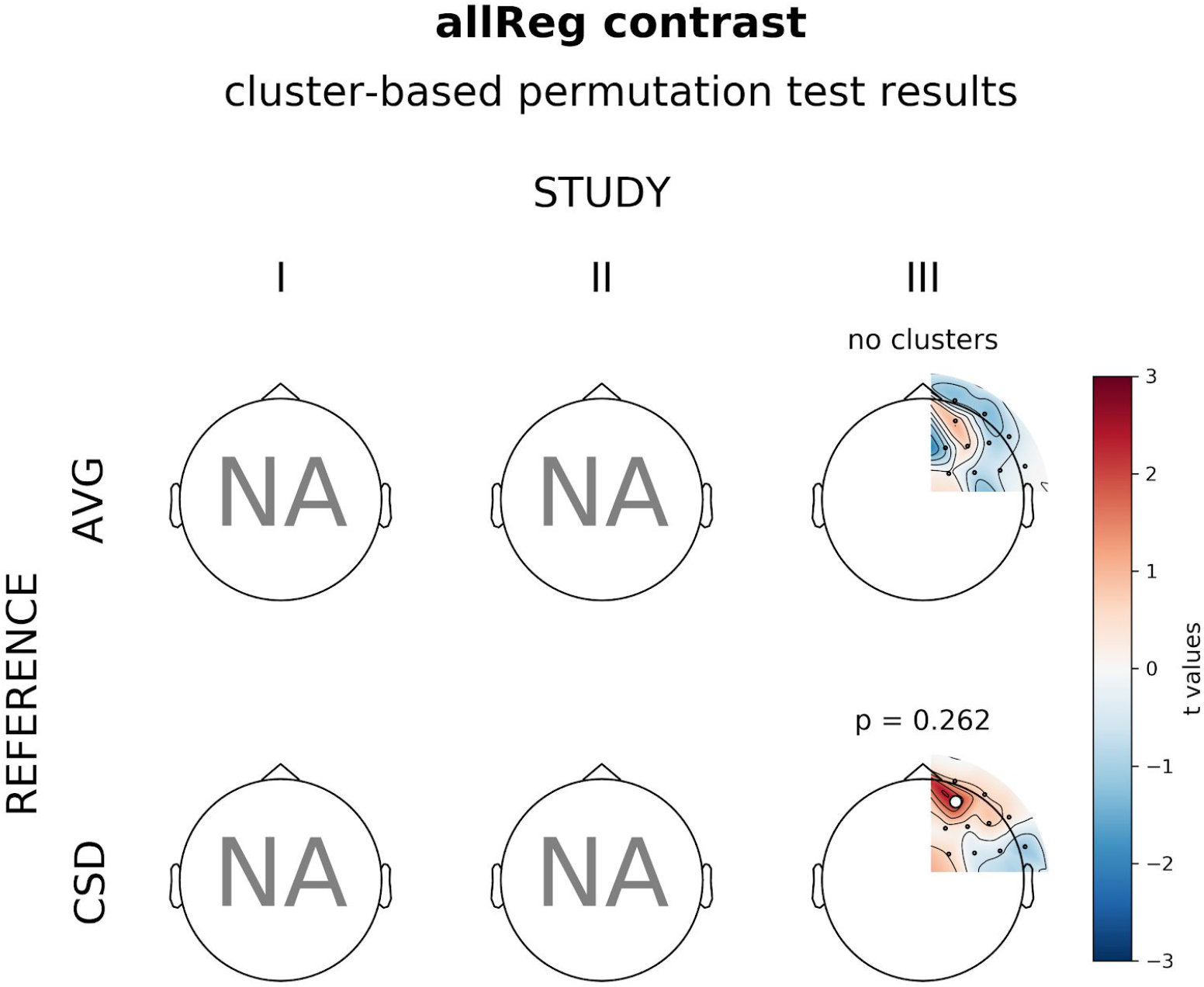
Results of cluster-based analyses for *allReg* contrast (regression on all subjects). Topographies of effects are presented in a reference by study matrix. Colorbar on the right presents color coding for the t values: more positive (red) / negative (blue) relationship between FAA and BDI. Channels that are part of a cluster are marked with white dots. NA means that the contrast was not available in given study.

**Figure 4 - figure supplement 2:**
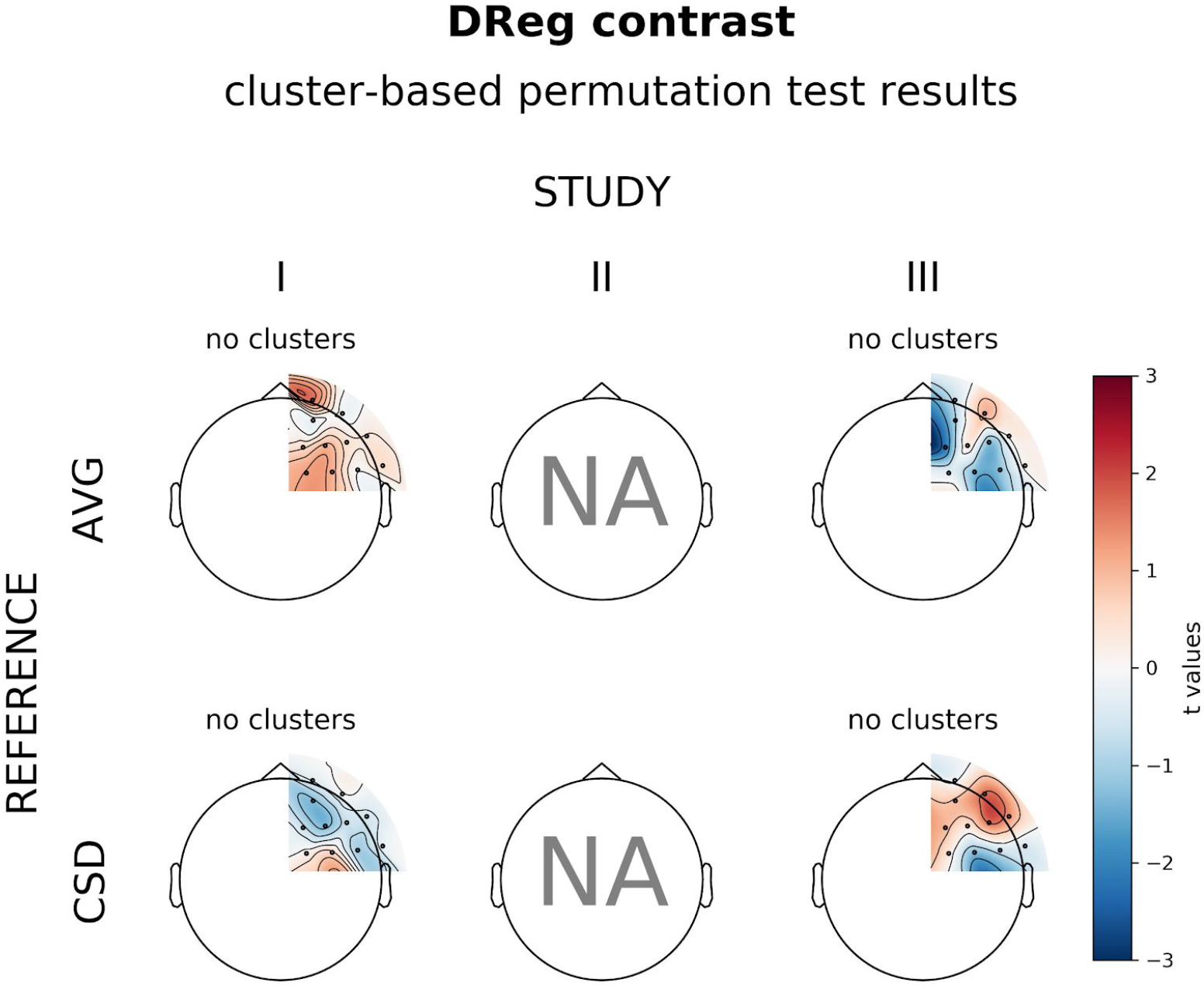
Results of cluster-based analyses for *DReg* contrast (regression on diagnosed subjects). Topographies of effects are presented in a reference by study matrix. Colorbar on the right presents color coding for the t values: more positive (red) / negative (blue) relationship between FAA and BDI. Channels that are part of a cluster are marked with white dots. NA means that the contrast was not available in given study.

**Figure 4 - figure supplement 3:**
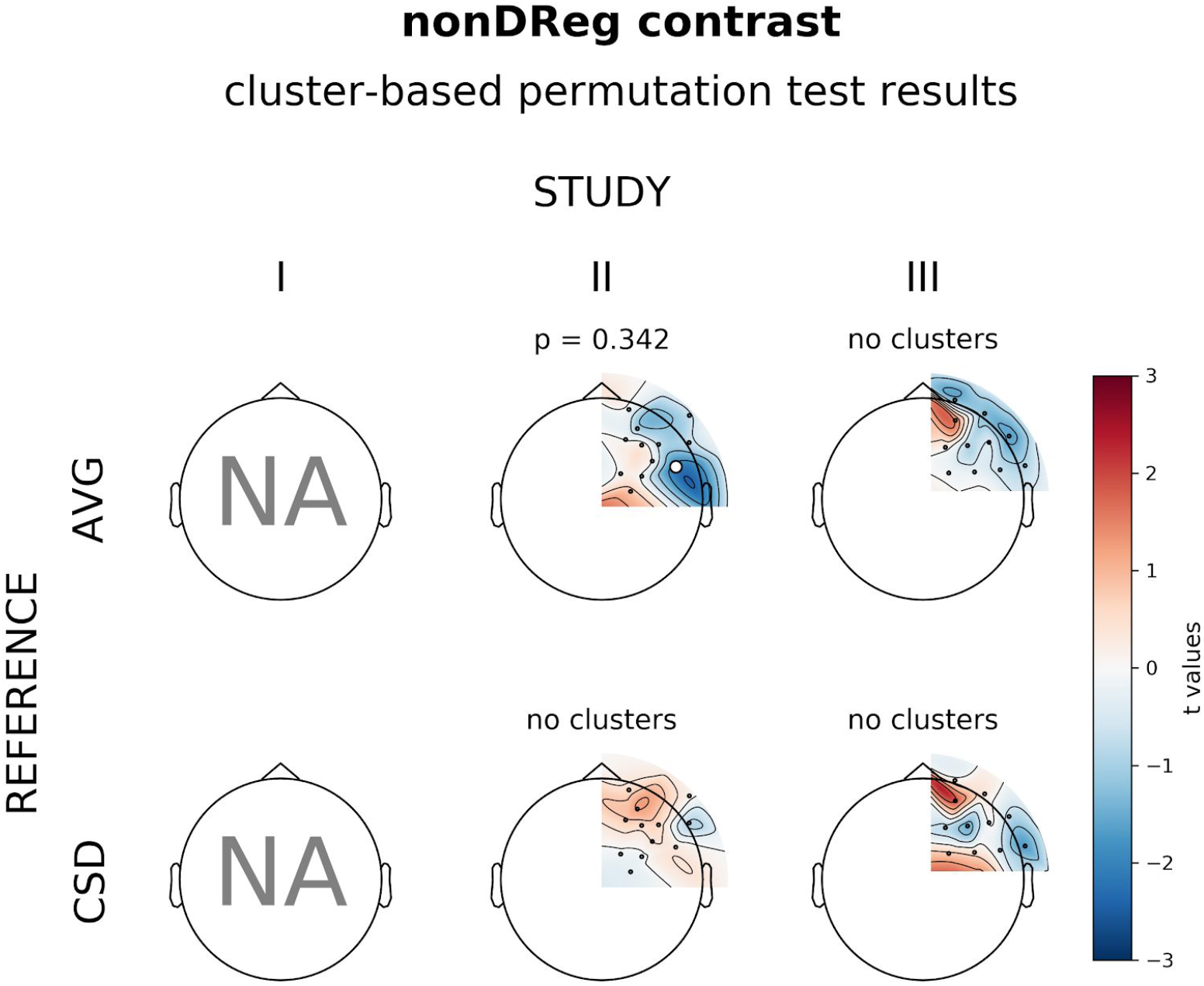
Results of cluster-based analyses for *nonDReg* contrast (regression on non diagnosed subjects). Topographies of effects are presented in a reference by study matrix. Colorbar on the right presents color coding for the t values: more positive (red) / negative (blue) relationship between FAA and BDI. Channels that are part of a cluster are marked with white dots. NA means that the contrast was not available in given study.

**Figure 4 - figure supplement 4:**
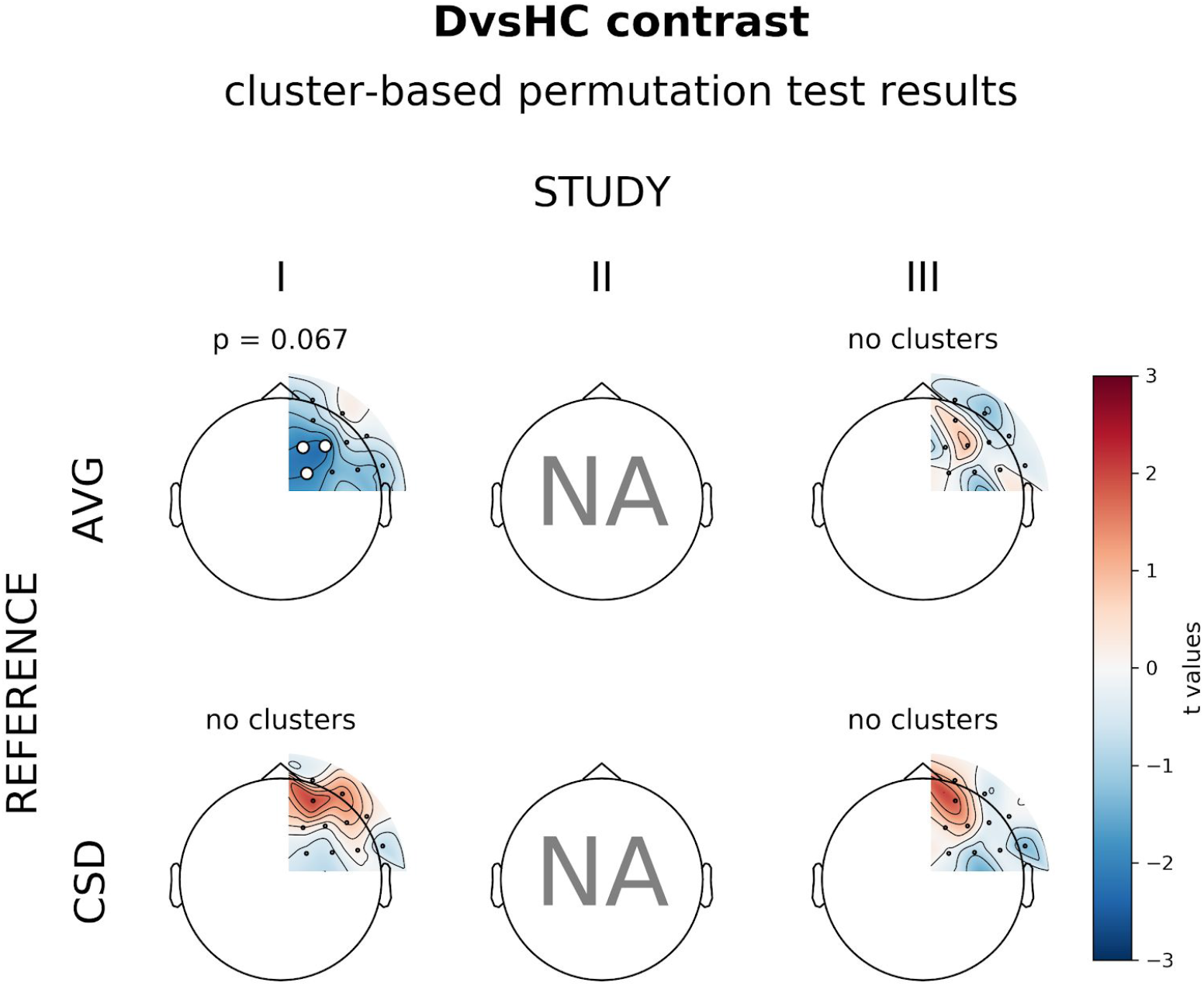
Results ofcluster-based analyses for *DvsHC* (comparison between diagnosed and healthy controls) contrast. Topographies of different contrast effects in a reference by study matrix. More positive (red) t values indicate more right-sided (less left-sided) alpha asymmetry for diagnosed participants. More negative (blue) t values indicate more right-sided (less left-sided) FAA for healthy controls. Channels that are part of a cluster are marked with white dots. NA means that the contrast was not available in given study.

**Figure 4 - figure supplement 5:**
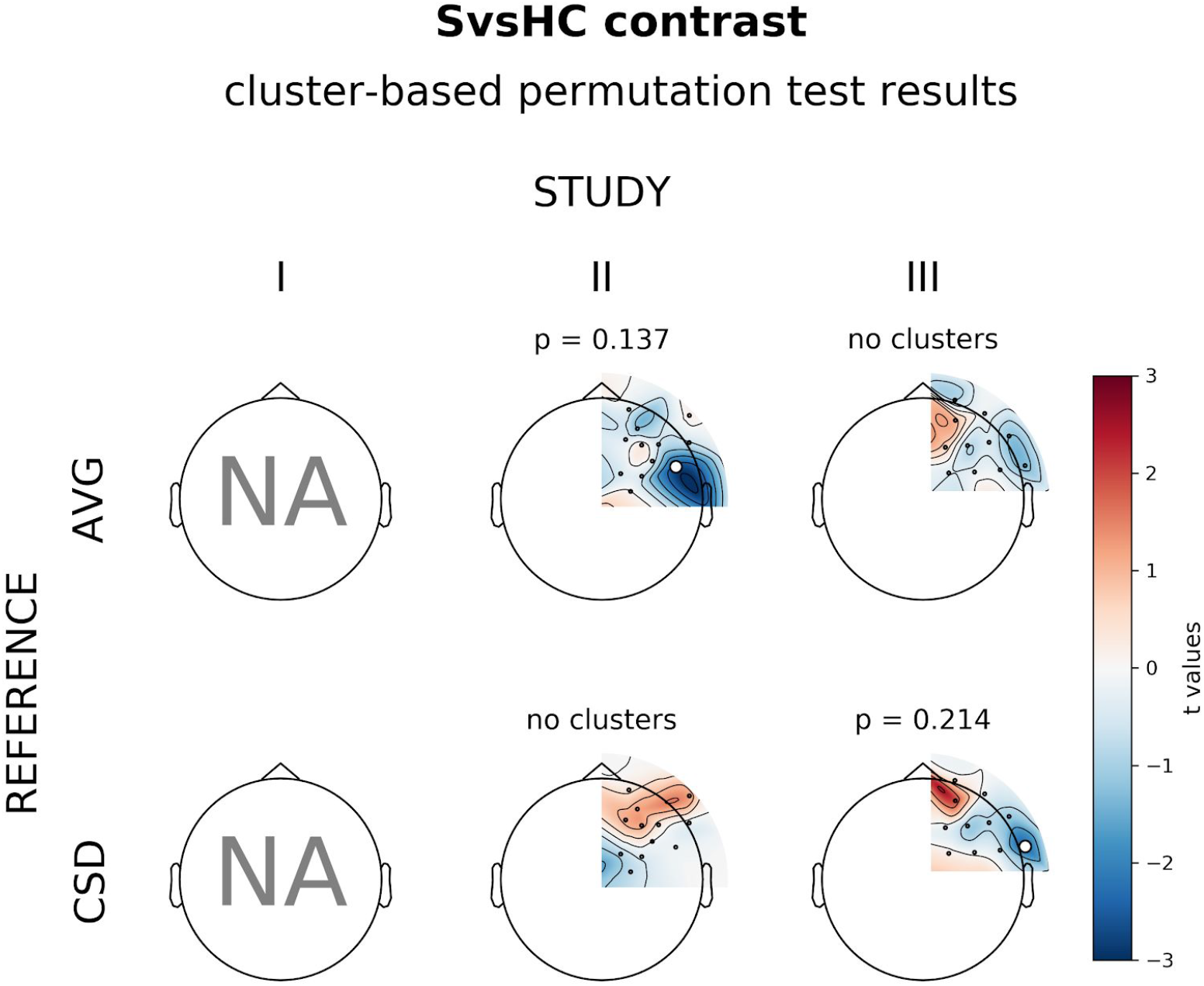
Results of cluster-based analyses for *SvsHC* (comparison between subclinical and healthy controls) contrast. Topographies of different contrast effects in a reference by study matrix. More positive (red) t values indicate more right-sided (less left-sided) alpha asymmetry for subclinical participants. More negative (blue) t values indicate more right-sided (less left-sided) FAA for healthy controls. Channels that are part of a cluster are marked with white dots. NA means that the contrast was not available in given study.

**Figure 5 - supplements 1-4: All results of cluster-based analyses on standardized data**

**Figure 5 - figure supplement 1:**
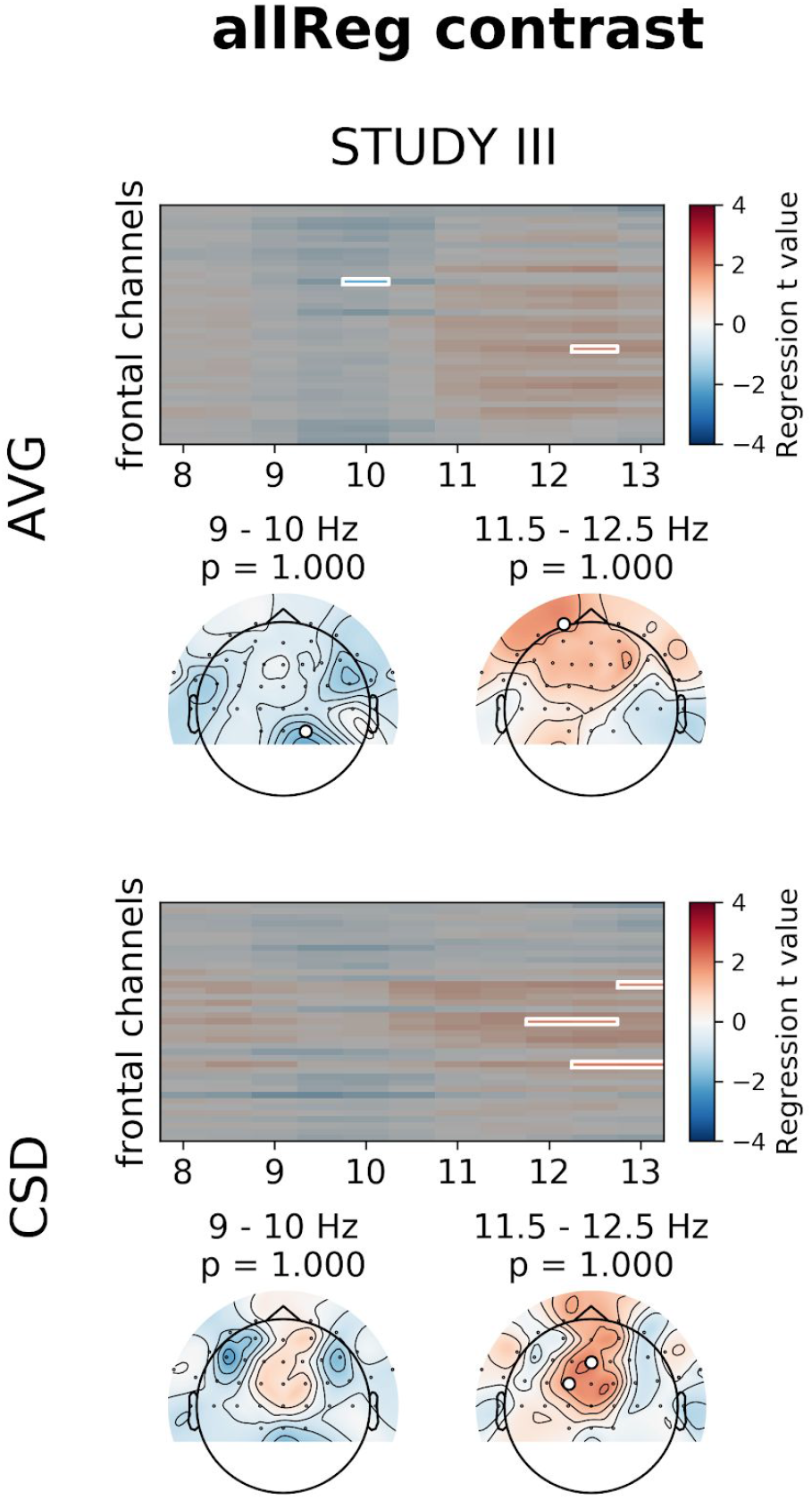
Results of cluster-based analyses on standardized data for *allReg* contrast (linear regression between FAA and BDI on all subjects together). Heatmaps in the upper part of each panel represent regression t-values for channel by frequency search space. More positive / negative t values, indicate higher / lower power with higher BDI. Clusters are indicated in the heatmaps with white outline. In each panel we present two topographies below the heatmap: showing average effect for 9 - 10 Hz and 11.5 - 12.5 Hz range respectively. Channels that are part of a cluster are marked with white dots in the topographical plots.

**Figure 5 - figure supplement 2:**
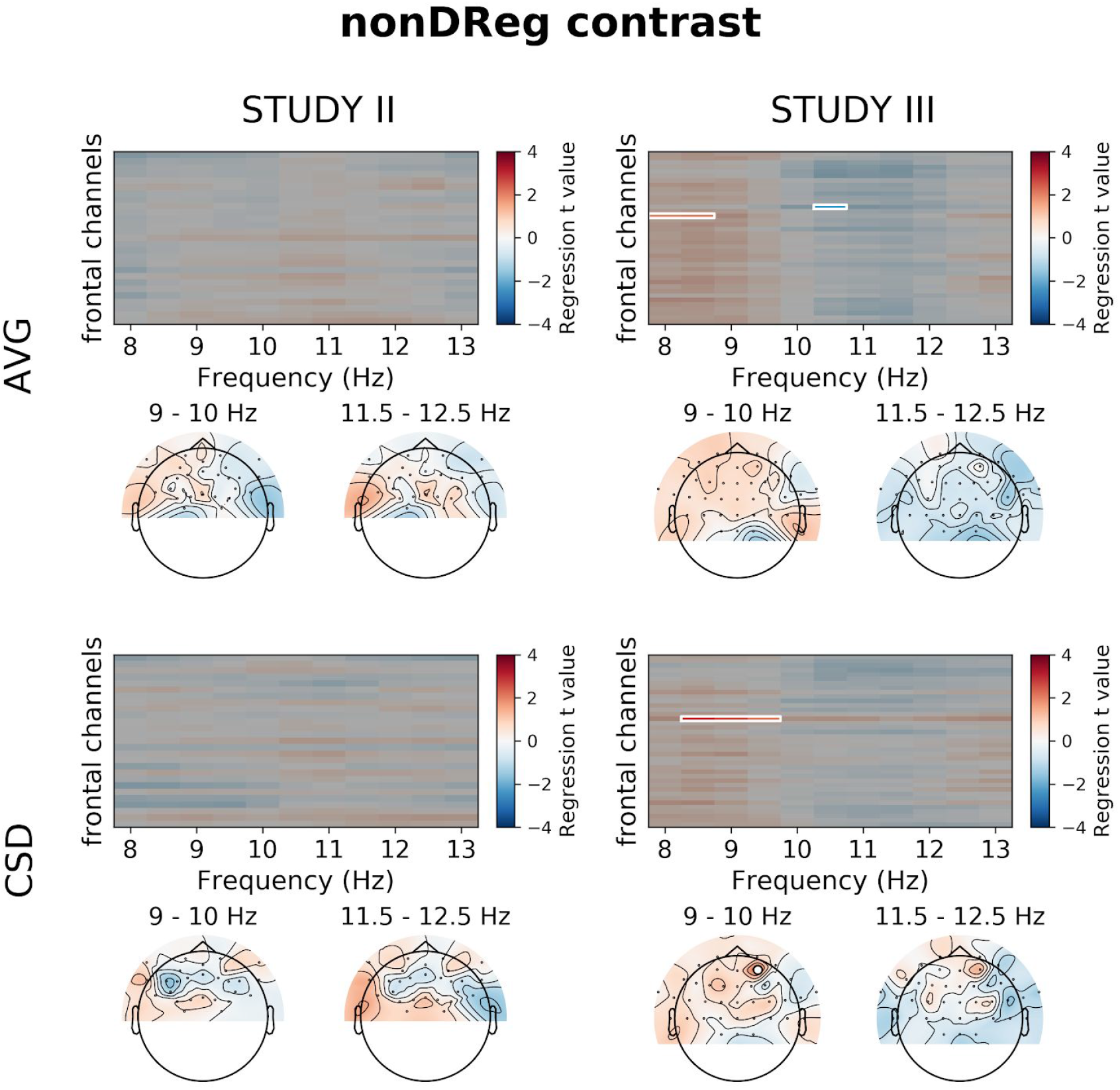
Results of cluster-based analyses on standardized data for *nonDReg* contrast (linear regression between FAA and BDI restricted to the non diagnosed subjects). Heatmaps in the upper part of each panel represent regression t-values for channel by frequency search space. More positive / negative t values, indicate higher / lower power with higher BDI. Clusters are indicated in the heatmaps with white outline. In each panel we present two topographies below the heatmap: showing average effect for 9 - 10 Hz and 11.5 - 12.5 Hz range respectively. Channels that are part of a cluster are marked with white dots in the topographical plots.

**Figure 5 - figure supplement 3:**
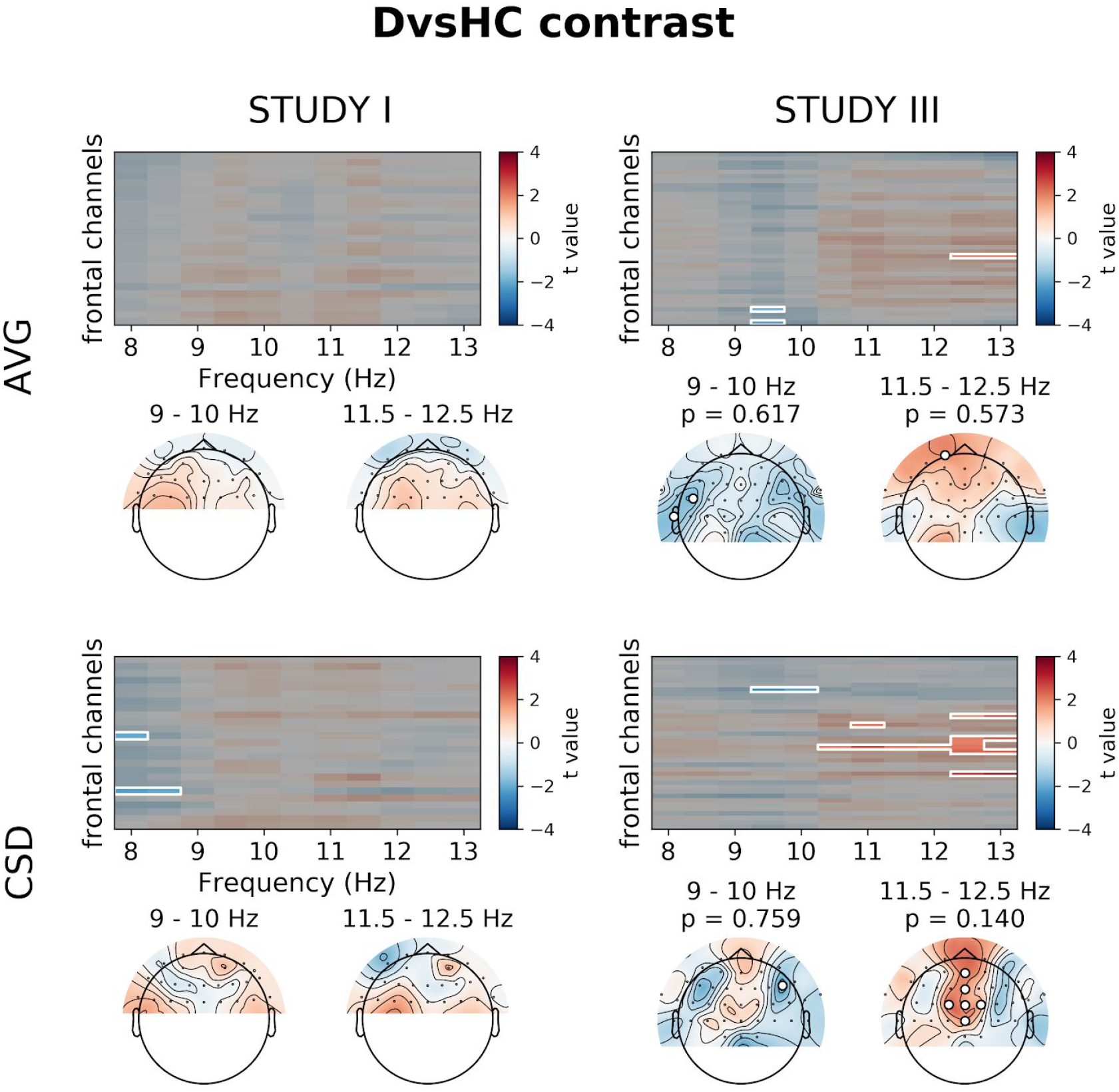
Results of cluster-based analyses on standardized data for *DvsHC* contrast (comparison between diagnosed and healthy controls). Heatmaps in the upper part of each panel represent t-values for channel by frequency search space. More positive (red) t values indicate higher power in given channel and frequency for diagnosed participants. More negative (blue) t values indicate higher power in given channel and frequency for healthy controls. Clusters are indicated in the heatmaps with white outline. In each panel we present two topographies below the heatmap: showing average effect for 9 - 10 Hz and 11.5 - 12.5 Hz range respectively. Channels that are part of a cluster are marked with white dots in the topographical plots.

**Figure 5 - figure supplement 4:**
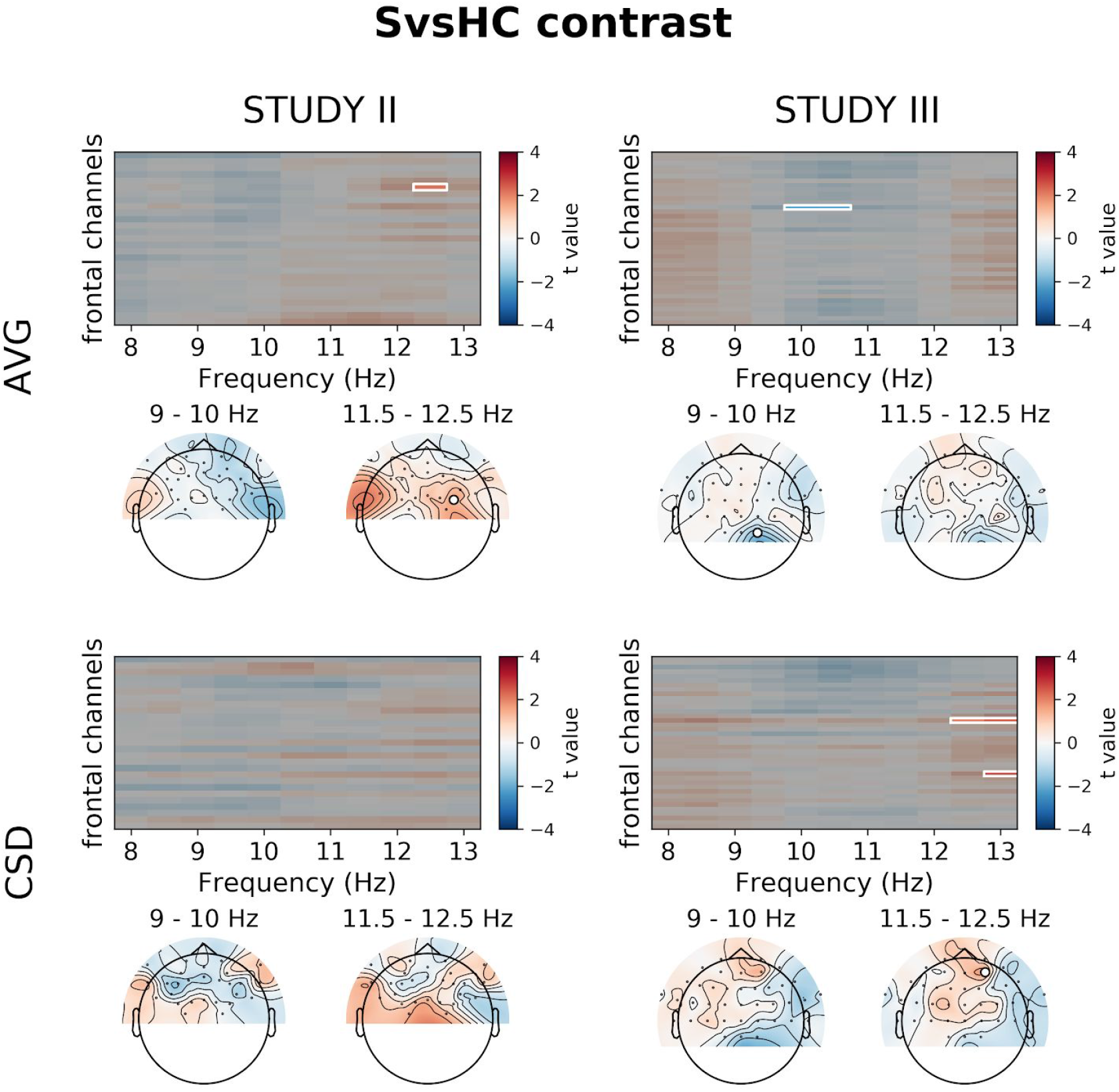
Results of cluster-based analyses on standardized data for *SvsHC* contrast (comparison between subclinical and healthy controls). Heatmaps in the upper part of each panel represent t-values for channel by frequency search space. More positive (red) t values indicate higher power in given channel and frequency for diagnosed participants. More negative (blue) t values indicate higher power in given channel and frequency for healthy controls. Clusters are indicated in the heatmaps with white outline. In each panel we present two topographies below the heatmap: showing average effect for 9 - 10 Hz and 11.5 - 12.5 Hz range respectively. Channels that are part of a cluster are marked with white dots in the topographical plots.

**Figure 6 - supplements 1-5: All results of source level analyses**

**Figure 6 - figure supplement 1:**
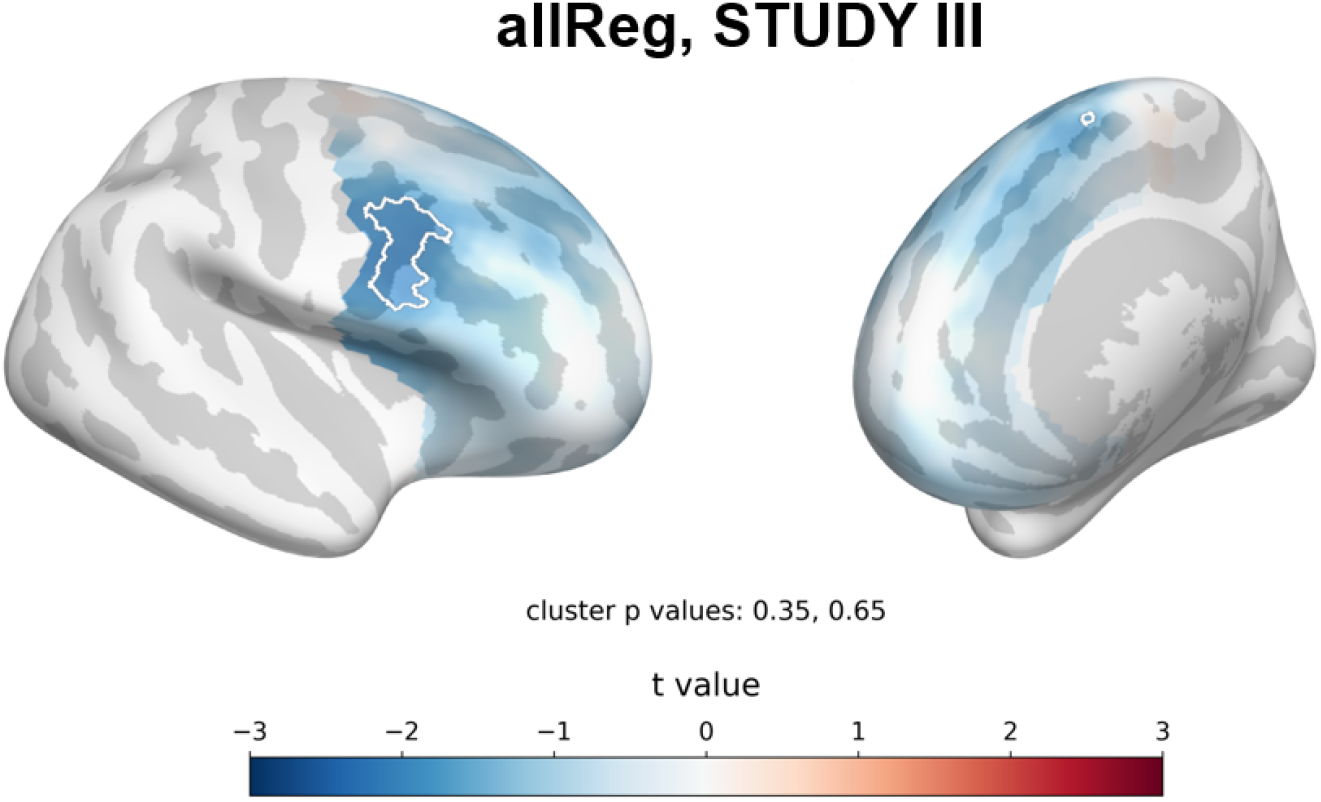
Results of source level analyses for *allReg* contrast (linear regression between FAA and BDI on all subjects) in Study III showing spatial t-value maps for regression analyses. Cluster limits are marked with white outlines, corresponding cluster p values are shown below each panel. Colorbar at the bottom presents color coding for the t values: more positive (red) / negative (blue) relationship between FAA and BDI. No cluster with p < 0.05 is present.

**Figure 6 - figure supplement 2:**
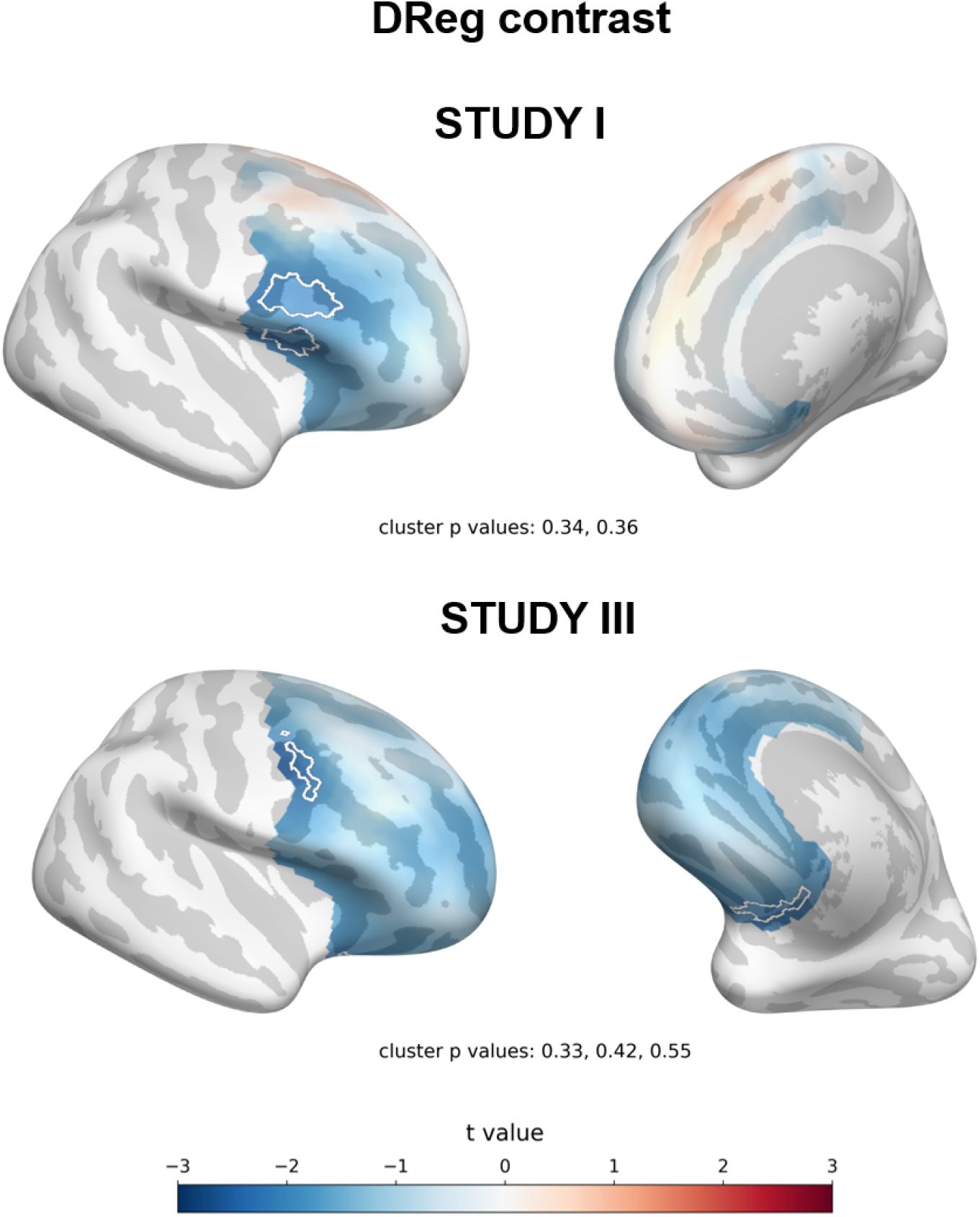
Results of source level analyses for *DReg* contrast (linear regression between FAA and BDI restricted to diagnosed subjects) in Study I and III showing spatial t-value maps for regression analyses. Cluster limits are marked with white outlines, corresponding cluster p values are shown below each panel. Colorbar at the bottom presents color coding for the t values: more positive (red) / negative (blue) relationship between FAA and BDI. Linear regression restricted to diagnosed subjects; no cluster with p < 0.05 is present.

**Figure 6 - figure supplement 3:**
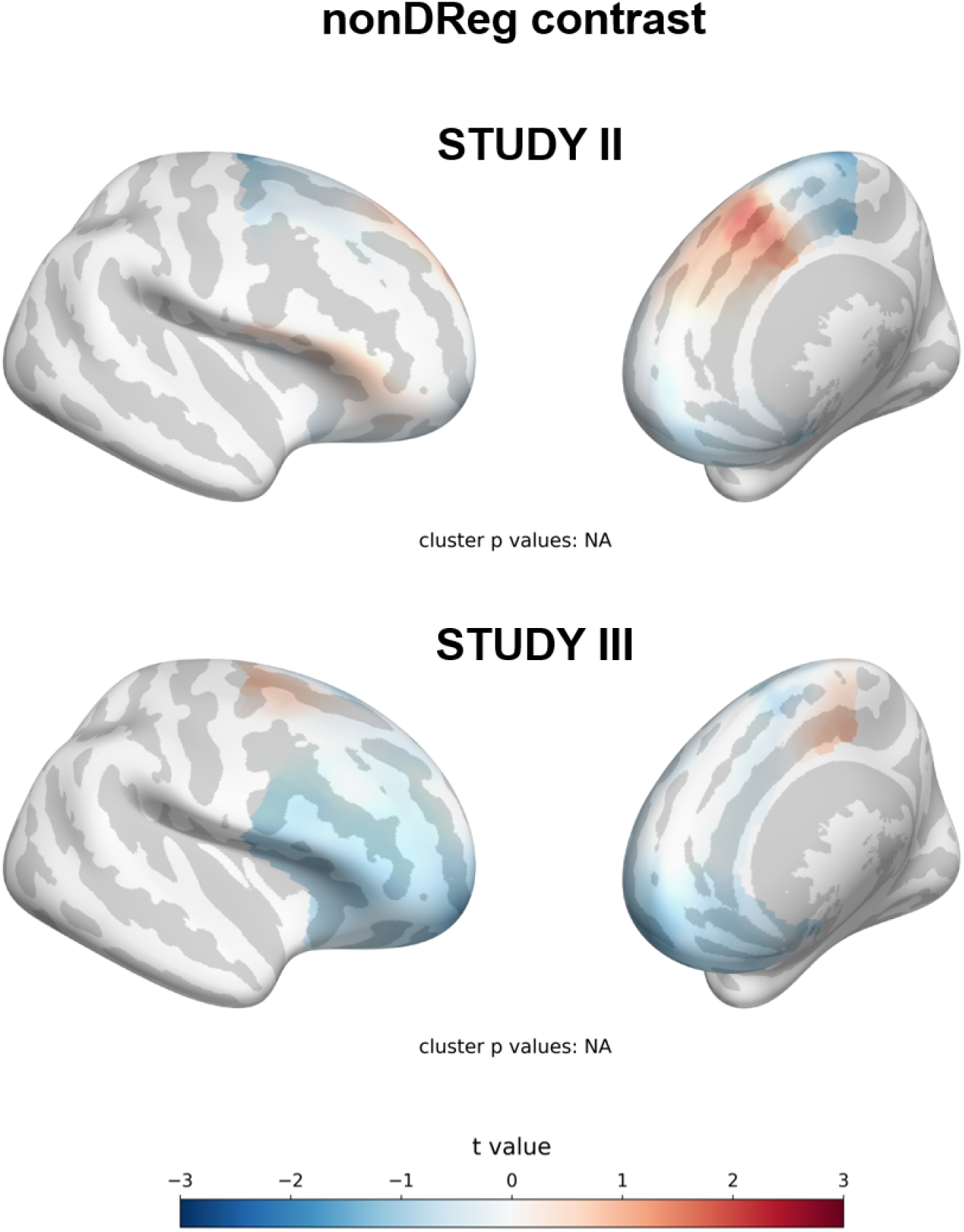
Results of source level analyses for *nonDReg* contrast (only the non diagnosed subjects) in Study II and III showing spatial t-value maps for regression analyses. Cluster limits are marked with white outlines, corresponding cluster p values are shown below each panel. Colorbar at the bottom presents color coding for the t values: more positive (red) / negative (blue) relationship between FAA and BDI. Linear regression restricted to all non diagnosed subjects; no cluster was found.

**Figure 6 - figure supplement 4:**
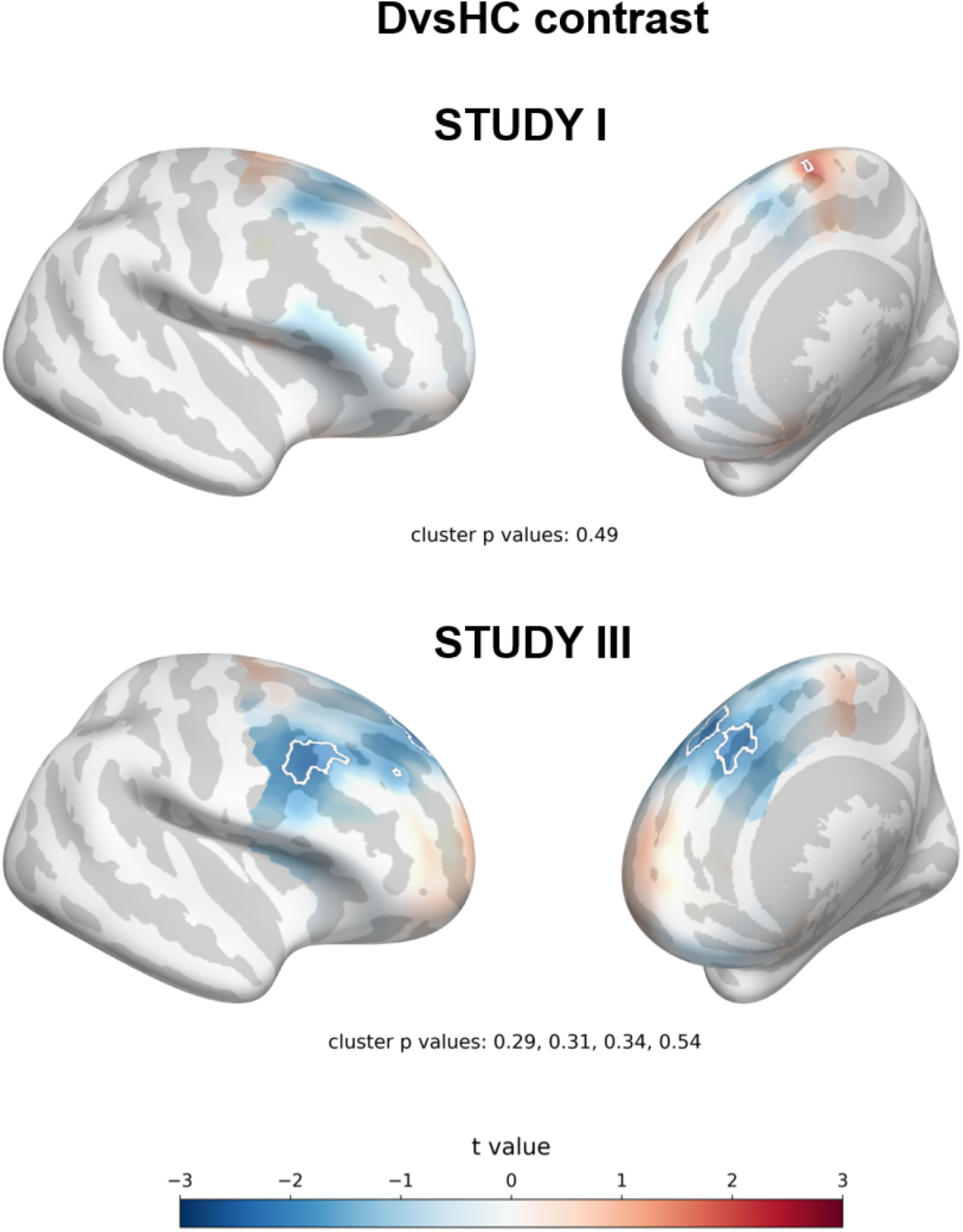
Results of source level analyses for *DvsHC* contrast (comparison between diagnosed and healthy controls) in Study I and III showing spatial t-value maps for regression analyses. Cluster limits are marked with white outlines, corresponding cluster p values are shown below each panel. More positive (red) t values indicate more right-sided (less left-sided) alpha asymmetry for diagnosed participants. More negative (blue) t values indicate more right-sided (less left-sided) FAA for healthy controls. No cluster with p < 0.05 is present.

**Figure 6 - figure supplement 5:**
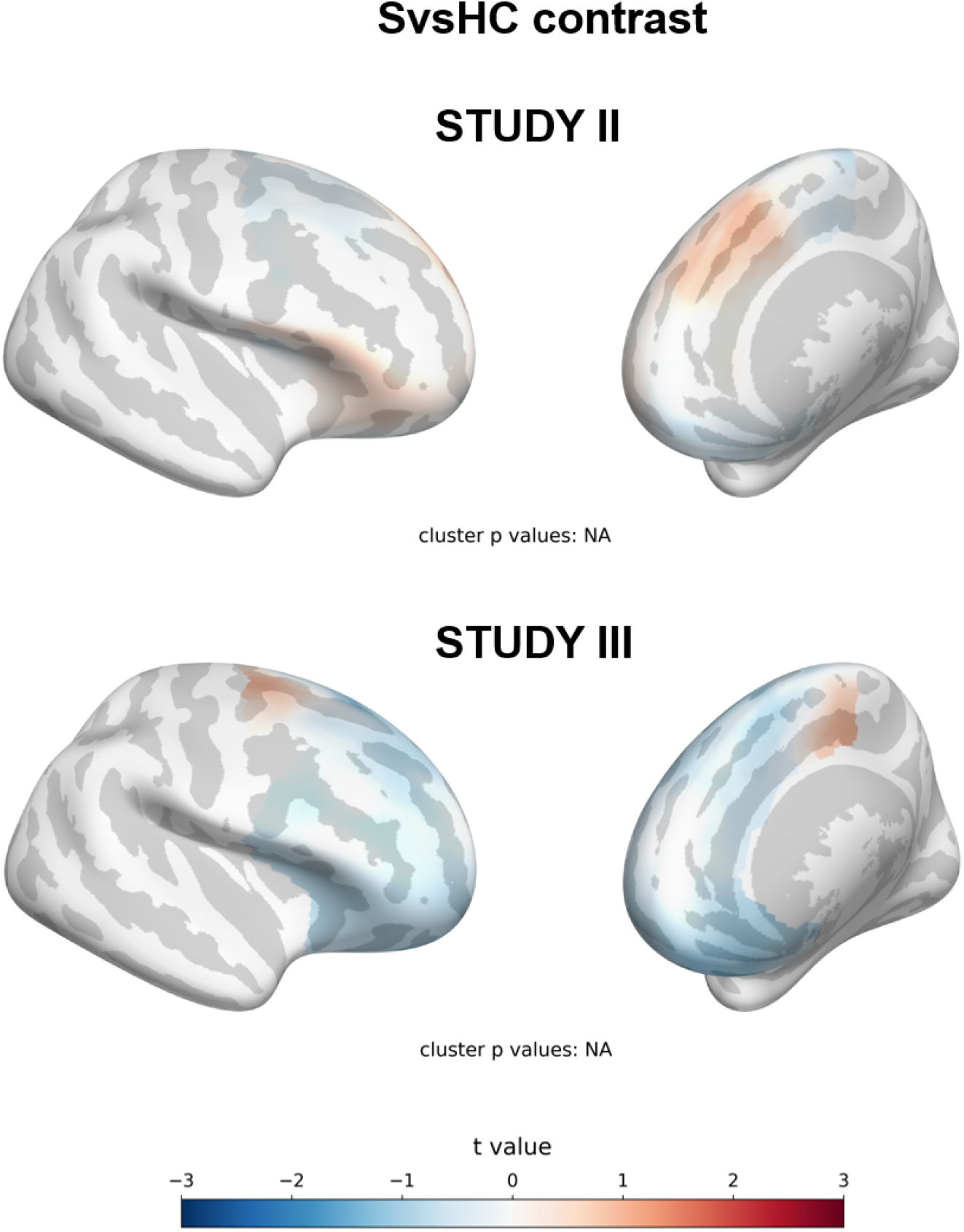
Results of source level analyses for *SvsHC* contrast (comparison between subclinical and healthy controls) in Study I and III showing spatial t-value maps for regression analyses. Cluster limits are marked with white outlines, corresponding cluster p values are shown below each panel. More positive (red) t values indicate more right-sided (less left-sided) alpha asymmetry for subclinical participants. More negative (blue) t values indicate more right-sided (less left-sided) FAA for healthy controls. No cluster was found.

## References

Allen, J. J. B., Urry, H. L., Hitt, S. K., & Coan, J. A. (2004). The stability of resting frontal electroencephalographic asymmetry in depression. Psychophysiology, 41(2), 269–280.

Arns, M., Bruder, G., Hegerl, U., Spooner, C., Palmer, D. M., Etkin, A., Fallahpour, K., Gatt, J. M., Hirshberg, L., & Gordon, E. (2016). EEG alpha asymmetry as a gender-specific predictor of outcome to acute treatment with different antidepressant medications in the randomized iSPOT-D study. Clinical Neurophysiology: Official Journal of the International Federation of Clinical Neurophysiology, 127(1), 509–519.

Baskaran, A., Milev, R., & McIntyre, R. S. (2012). The neurobiology of the EEG biomarker as a predictor of treatment response in depression. Neuropharmacology, 63(4), 507–513.

Beck, A., Steer, R., & Brown, G. (1996). Manual for the Beck depression inventory-II (BDI-II). https://www.scienceopen.com/document?vid=9feb932d-1f91-4ff9-9d27-da3bda716129

Beck, A. T., Ward, C. H., Mendelson, M., Mock, J., & Erbaugh, J. (1961). An inventory for measuring depression. Archives of General Psychiatry, 4, 561–571.

Beeney, J. E., Levy, K. N., Gatzke-Kopp, L. M., & Hallquist, M. N. (2014). EEG asymmetry in borderline personality disorder and depression following rejection. In Personality Disorders: Theory, Research, and Treatment (Vol. 5, Issue 2, pp. 178–185). https://doi.org/10.1037/per0000032

Botvinik-Nezer, R., Holzmeister, F., Camerer, C. F., Dreber, A., Huber, J., Johannesson, M., Kirchler, M., Iwanir, R., Mumford, J. A., Adcock, A., & Others. (2019). Variability in the analysis of a single neuroimaging dataset by many teams. BioRxiv, 843193.

Carvalho, A., Moraes, H., Silveira, H., Ribeiro, P., Piedade, R. A. M., Deslandes, A. C., Laks, J., & Versiani, M. (2011). EEG frontal asymmetry in the depressed and remitted elderly: Is it related to the trait or to the state of depression? Journal of Affective Disorders, 129(1-3), 143–148.

Cohen, M. X. (2015). Comparison of different spatial transformations applied to EEG data: A case study of error processing. International Journal of Psychophysiology: Official Journal of the International Organization of Psychophysiology, 97(3), 245–257.

Cohen, M. X., & Gulbinaite, R. (2014). Five methodological challenges in cognitive electrophysiology. NeuroImage, 85 Pt 2, 702–710.

Dale, A. M., Fischl, B., & Sereno, M. I. (1999). Cortical surface-based analysis. I. Segmentation and surface reconstruction. NeuroImage, 9(2), 179–194.

Davidson, R. J. (1979). Frontal Versus Perietal EEG Asymmetry during Positive and Negative Affect. Psychophysiology, 16(2), 202–203.

Davidson, R. J. (1984). 11 Affect, cognition, and hemispheric specialization. Emotions, Cognition, and Behavior, 320–365.

Davidson, R. J. (2004). What does the prefrontal cortex “do” in affect: perspectives on frontal EEG asymmetry research. Biological Psychology, 67(1), 219–234.

de Aguiar Neto, F. S., & Rosa, J. L. G. (2019). Depression biomarkers using non-invasive EEG: A review. Neuroscience and Biobehavioral Reviews, 105, 83–93.

Deldin, P. J., & Chiu, P. (2005). Cognitive restructuring and EEG in major depression. Biological Psychology, 70(3), 141–151.

Delorme, A., & Makeig, S. (2004). EEGLAB: an open source toolbox for analysis of single-trial EEG dynamics including independent component analysis. Journal of Neuroscience Methods, 134(1), 9–21.

Fischl, B., Sereno, M. I., Tootell, R. B., & Dale, A. M. (1999). High-resolution intersubject averaging and a coordinate system for the cortical surface. Human Brain Mapping, 8(4), 272–284.

Gelman, A., & Loken, E. (2014). The Statistical Crisis in Science. In American Scientist (Vol. 102, Issue 6, p. 460). https://doi.org/10.1511/2014.111.460

Gold, C., Fachner, J., & Erkkilä, J. (2013). Validity and reliability of electroencephalographic frontal alpha asymmetry and frontal midline theta as biomarkers for depression. In Scandinavian Journal of Psychology (Vol. 54, Issue 2, pp. 118–126). https://doi.org/10.1111/sjop.12022

Gramfort, A., Luessi, M., Larson, E., Engemann, D. A., Strohmeier, D., Brodbeck, C., Goj, R., Jas, M., Brooks, T., Parkkonen, L., & Hämäläinen, M. (2013). MEG and EEG data analysis with MNE-Python. Frontiers in Neuroscience, 7, 267.

Gramfort, A., Luessi, M., Larson, E., Engemann, D. A., Strohmeier, D., Brodbeck, C., Parkkonen, L., & Hämäläinen, M. S. (2014). MNE software for processing MEG and EEG data. NeuroImage, 86, 446–460.

Greve, D. N., Van der Haegen, L., Cai, Q., Stufflebeam, S., Sabuncu, M. R., Fischl, B., & Brysbaert, M. (2013). A surface-based analysis of language lateralization and cortical asymmetry. Journal of Cognitive Neuroscience, 25(9), 1477–1492.

Gross, J., Kujala, J., Hamalainen, M., Timmermann, L., Schnitzler, A., & Salmelin, R. (2001). Dynamic imaging of coherent sources: Studying neural interactions in the human brain. Proceedings of the National Academy of Sciences of the United States of America, 98(2), 694–699.

Hipp, J. F., & Siegel, M. (2013). Dissociating neuronal gamma-band activity from cranial and ocular muscle activity in EEG. In Frontiers in Human Neuroscience (Vol. 7). https://doi.org/10.3389/fnhum.2013.00338

Ho, J., Tumkaya, T., Aryal, S., Choi, H., & Claridge-Chang, A. (2019). Moving beyond P values: data analysis with estimation graphics. Nature methods, 16(7), 565–566.

Iosifescu, D. V., Greenwald, S., Devlin, P., Mischoulon, D., Denninger, J. W., Alpert, J. E., & Fava, M. (2009). Frontal EEG predictors of treatment outcome in major depressive disorder. European Neuropsychopharmacology: The Journal of the European College of Neuropsychopharmacology, 19(11), 772–777.

Jiang, H., Popov, T., Jylänki, P., Bi, K., Yao, Z., Lu, Q., Jensen, O., & van Gerven, M. A. J. (2016). Predictability of depression severity based on posterior alpha oscillations. Clinical Neurophysiology: Official Journal of the International Federation of Clinical Neurophysiology, 127(4), 2108–2114.

Kaiser, A., Doppelmayr, M., & Iglseder, B. (2018). Electroencephalogram alpha asymmetry in geriatric depression. Zeitschrift Fur Gerontologie Und Geriatrie, 51(2), 200–205.

Kaiser, A. K., Gnjezda, M.-T., Knasmüller, S., & Aichhorn, W. (2018). Electroencephalogram alpha asymmetry in patients with depressive disorders: current perspectives. Neuropsychiatric Disease and Treatment, 14, 1493–1504.

Kemp, A. H., Griffiths, K., Felmingham, K. L., Shankman, S. A., Drinkenburg, W., Arns, M., Clark, C. R., & Bryant, R. A. (2010). Disorder specificity despite comorbidity: resting EEG alpha asymmetry in major depressive disorder and post-traumatic stress disorder. Biological Psychology, 85(2), 350–354.

Kentgen, L. M., Tenke, C. E., Pine, D. S., Fong, R., Klein, R. G., & Bruder, G. E. (2000). Electroencephalographic asymmetries in adolescents with major depression: Influence of comorbidity with anxiety disorders. In Journal of Abnormal Psychology (Vol. 109, Issue 4, pp. 797–802). https://doi.org/10.1037/0021-843x.109.4.797

Knott, V., Mahoney, C., Kennedy, S., & Evans, K. (2001). EEG power, frequency, asymmetry and coherence in male depression. Psychiatry Research, 106(2), 123–140.

Koles, Z. J., Lazar, M. S., & Zhou, S. Z. (1990). Spatial patterns underlying population differences in the background EEG. Brain Topography, 2(4), 275–284.

Lubar, J. F., Congedo, M., & Askew, J. H. (2003). Low-resolution electromagnetic tomography (LORETA) of cerebral activity in chronic depressive disorder. In International Journal of Psychophysiology (Vol. 49, Issue 3, pp. 175–185). https://doi.org/10.1016/s0167-8760(03)00115-6

Magnuski, M. (2020a). GitHub repository: borsar. Borsar. https://github.com/mmagnuski/borsar

Magnuski, M. (2020b). GitHub repository: eegDb. eegDb. https://github.com/mmagnuski/eegDb

Magnuski, M. (2020c). GitHub repository: sarna. Sarna. https://github.com/mmagnuski/sarna

Magnuski, M., & Ruban, A. (2020). GitHub repository: DiamSar. DiamSar. https://github.com/mmagnuski/DiamSar

Maris, E., & Oostenveld, R. (2007). Nonparametric statistical testing of EEG- and MEG-data. Journal of Neuroscience Methods, 164(1), 177–190.

Mathersul, D., Williams, L. M., Hopkinson, P. J., & Kemp, A. H. (2008). Investigating models of affect: relationships among EEG alpha asymmetry, depression, and anxiety. Emotion, 8(4), 560–572.

McMenamin, B. W., Shackman, A. J., Maxwell, J. S., Bachhuber, D. R. W., Koppenhaver, A. M., Greischar, L. L., & Davidson, R. J. (2010). Validation of ICA-based myogenic artifact correction for scalp and source-localized EEG. NeuroImage, 49(3), 2416–2432.

Oostenveld, R., Fries, P., Maris, E., & Schoffelen, J.-M. (2011). FieldTrip: Open source software for advanced analysis of MEG, EEG, and invasive electrophysiological data. Computational Intelligence and Neuroscience, 2011, 156869.

Parra, L., & Sajda, P. (2003). Blind Source Separation via Generalized Eigenvalue Decomposition. Journal of Machine Learning Research: JMLR, 4(Dec), 1261–1269.

Schaffer, C. E., Davidson, R. J., & Saron, C. (1983). Frontal and parietal electroencephalogram asymmetry in depressed and nondepressed subjects. Biological Psychiatry, 18(7), 753–762.

Shackman, A. J., McMenamin, B. W., Slagter, H. A., Maxwell, J. S., Greischar, L. L., & Davidson, R. J. (2009). Electromyogenic artifacts and electroencephalographic inferences. Brain Topography, 22(1), 7–12.

Sheehan, D. V., Lecrubier, Y., Sheehan, K. H., Amorim, P., Janavs, J., Weiller, E., Hergueta, T., Baker, R., & Dunbar, G. C. (1998). The Mini-International Neuropsychiatric Interview (MINI): the development and validation of a structured diagnostic psychiatric interview for DSM-IV and ICD-10. The Journal of Clinical Psychiatry. https://psycnet.apa.org/record/1998-03251-004

Smith, E. E., Cavanagh, J. F., & Allen, J. J. B. (2018). Intracranial source activity (eLORETA) related to scalp-level asymmetry scores and depression status. In Psychophysiology (Vol. 55, Issue 1, p. e13019). https://doi.org/10.1111/psyp.13019

Smith, E. E., Reznik, S. J., Stewart, J. L., & Allen, J. J. B. (2017). Assessing and conceptualizing frontal EEG asymmetry: An updated primer on recording, processing, analyzing, and interpreting frontal alpha asymmetry. International Journal of Psychophysiology: Official Journal of the International Organization of Psychophysiology, 111, 98–114.

Steegen, S., Tuerlinckx, F., Gelman, A., & Vanpaemel, W. (2016). Increasing Transparency Through a Multiverse Analysis. Perspectives on Psychological Science: A Journal of the Association for Psychological Science, 11(5), 702–712.

Stewart, J. L., Coan, J. A., Towers, D. N., & Allen, J. J. B. (2014). Resting and task-elicited prefrontal EEG alpha asymmetry in depression: support for the capability model. Psychophysiology, 51(5), 446–455.

Szumska, I., Gola, M., Rusanowska, M., Łempicka, M., Żygierewicz, J., Krejtz, I., Nezlek, J. B., & Holas, P. (2020). Mindfulness-based cognitive therapy reduces clinical symptoms, but do not change frontal alpha asymmetry in people with major depression disorder, https://doi.org/10.31234/osf.io/4bzgs

Tadel, F., Baillet, S., Mosher, J. C., Pantazis, D., & Leahy, R. M. (2011). Brainstorm: a user-friendly application for MEG/EEG analysis. Computational Intelligence and Neuroscience, 2011, 879716.

Thibodeau, R., Jorgensen, R. S., & Kim, S. (2006). Depression, anxiety, and resting frontal EEG asymmetry: a meta-analytic review. Journal of Abnormal Psychology, 115(4), 715–729.

Tibshirani, R. J., & Efron, B. (1993). An introduction to the bootstrap. Monographs on statistics and applied probability, 57, 1–436.

Toldo, R., Gherardi, R., Farenzena, M., & Fusiello, A. (2015). Hierarchical structure-and-motion recovery from uncalibrated images. Computer Vision and Image Understanding: CVIU, 140, 127–143.

Tomé, A. M. (2006). The generalized eigendecomposition approach to the blind source separation problem. Digital Signal Processing, 16(3), 288–302.

van der Meij, R., Jacobs, J., & Maris, E. (2015). Uncovering phase - coupled oscillatory networks in electrophysiological data. Human Brain Mapping. https://onlinelibrary.wiley.com/doi/abs/10.1002/hbm.22798

van der Meij, R., van Ede, F., & Maris, E. (2016). Rhythmic Components in Extracranial Brain Signals Reveal Multifaceted Task Modulation of Overlapping Neuronal Activity. PloS One, 11(6), e0154881.

van der Vinne, N., Madelon A., V., Michel JAM, V. P., & Martijn, A. (2017). Frontal alpha asymmetry as a diagnostic marker in depression: Fact or fiction? A meta-analysis. In NeuroImage: Clinical (Vol. 16, pp. 79–87). https://doi.org/10.1016/j.nicl.2017.07.006

van Ede, F., & Maris, E. (2016). Physiological Plausibility Can Increase Reproducibility in Cognitive Neuroscience. Trends in Cognitive Sciences, 20(8), 567–569.

Van Veen, B. D., van Drongelen, W., Yuchtman, M., & Suzuki, A. (1997). Localization of brain electrical activity via linearly constrained minimum variance spatial filtering. IEEE Transactions on Bio-Medical Engineering, 44 (9), 867–880.

van Vliet, M., Liljeström, M., Aro, S., Salmelin, R., & Kujala, J. (2018). Analysis of Functional Connectivity and Oscillatory Power Using DICS: From Raw MEG Data to Group-Level Statistics in Python. Frontiers in Neuroscience, 12, 586.

Vuga, M., Fox, N. A., Cohn, J. F., George, C. J., Levenstein, R. M., & Kovacs, M. (2006). Long-term stability of frontal electroencephalographic asymmetry in adults with a history of depression and controls. International Journal of Psychophysiology: Official Journal of the International Organization of Psychophysiology, 59(2), 107–115.

## References

Rouder, J. N., Speckman, P. L., Sun, D., Morey, R. D., & Iverson, G. (2009). Bayesian t tests for accepting and rejecting the null hypothesis. Psychonomic bulletin & review, 16(2), 225–237.

Vallat, R. (2018). Pingouin: statistics in Python. Journal of Open Source Software, 3(31), 1026, https://doi.org/10.21105/joss.01026

